# AMBERff at scale: Multimillion-atom simulations with AMBER force fields in NAMD

**DOI:** 10.1101/2023.10.10.561755

**Authors:** Santiago Antolínez, Peter Eugene Jones, James C. Phillips, Jodi A. Hadden-Perilla

## Abstract

All-atom molecular dynamics (MD) simulations are an essential structural biology technique with increasing application to multimillion-atom systems, including viruses and cellular machinery. Classical MD simulations rely on parameter sets, such as the AMBER family of force fields (AMBERff), to accurately describe molecular motion. Here, we present an implementation of AMBERff for use in NAMD that overcomes previous limitations to enable high-performance, massively-parallel simulations encompassing up to two billion atoms. Single-point potential energy comparisons and case studies on model systems demonstrate that the implementation produces results that are as accurate as running AMBERff in its native engine.

## Introduction

All-atom molecular dynamics (MD) simulations have emerged as an essential structural biology technique, capable of revealing details that are inaccessible to experimental methods. When applied to large-scale biomolecular systems like viruses and cellular machinery, MD simulations serve as a computational microscope^1^ that enables researchers to investigate limited timescales of biological activity. A multitude of groundbreaking discoveries have been driven by large-scale all-atom MD simulations in recent years, particularly through the integration of experimental data from NMR spectroscopy, X-ray crystallography, and cryo-electron microscopy and tomography.^2,3^

Classical MD simulations model the atoms and bonds of biomolecular systems as balls and springs and rely on force fields to propagate their motion. A force field is a mathematical means to calculate the potential energy of a molecule as a function of its conformation (e.g., Eq. 1 and 2). From the gradient of potential energy, the forces acting on constituent atoms are determined and applied to drive configurational changes of the system over discrete simulation timesteps. By calculating millions upon millions of steps through time, sampling the conformational evolution of the biomolecular system over many moments, a trajectory of molecular motion is generated. Force fields are painstakingly parameterized, empirically optimized, and experimentally validated to capture realistic dynamics and reproduce biophysical properties. Force field development represents an active area of ongoing research.^4–6^ Force field families founded on self-consistent philosophies include parameter sets for distinct classes of biomolecules, which may be combined to simulate complex, chemically authentic systems. Widely used biomolecular force field families include AMBERff, CHARMMff, GROMOS, and OPLS.

MD simulation software are computational engines that apply force fields to biomolecular systems to model their motion. Major software packages for biomolecular simulation include AMBER,^7,8^ CHARMM,^9^ GROMACS,^10^ LAMMPS,^11^ and NAMD. ^12,13^ While AMBER’s *pmemd*.*cuda* ^14,15^ excels for small-scale simulations performed on single compute nodes with GPUs (biomolecular systems comprising tens to hundreds of thousands of atoms), NAMD2 excels for large-scale simulations performed on leadership-class supercomputers (biomolecular systems comprising millions to billions of atoms). NAMD3 is competitive on single, GPU-accelerated nodes for intermediate systems of up to several million atoms.

Importantly, at the multimillion-atom level, systems of true biological significance can be investigated, such as those probing macromolecular assemblies responsible for viral and cellular processes. The largest all-atom MD simulations reported have been carried out with CHARMMff in NAMD2, including the HIV-1 capsid (64 million atoms), ^16,17^ the photosynthetic chromatophore organelle (100 million atoms),^18^ and the influenza A virion (200 million atoms).^19^ Systems of this size are not feasible to build within the AMBER framework due to limitations in the fixed-column-width PRMTOP file format and the need to pre-calculate values for the PRMTOP using AMBER’s single-threaded file builder *tleap*.

Given increasing efforts in the field to model and simulate large biological systems, including intact viruses and minimal cells with their enclosed genetic material, the availability of all leading biomolecular force fields within software well-suited for high-performance simulations at scale is essential. Here, we present an implementation of AMBERff for use in NAMD, enabling simulations with the force field family encompassing up to two billion atoms — an increase of system size by three orders of magnitude beyond what is currently tractable with AMBERff in its native engine. Our approach is straightforward, builds upon existing software infrastructure, and produces excellent potential energy agreement, while accurately reproducing biophysical properties. The implementation relies on the *psfgen* file builder, available as both a VMD^20^ plugin and a stand-alone binary distributed with NAMD, to generate PSF and JS molecular topology files that are not constrained by atom count, allowing AMBERff to be applied to study the dynamics of biological systems at scale.

## Methods

### Approach

Molecular topology and parameter information for AMBERff are encoded in the PRMTOP file format (produced by e.g., *tleap*), which is read natively by the AMBER software. PRMTOP files can be read directly by NAMD, already allowing accurate simulations with the force field on supercomputers (used for example in Ref. ^21^). However, because the AMBER software is not designed for large-scale simulations, PRMTOP files with their fixed column widths do not accommodate large atom counts. Eight digits are allocated for any single integer contained in the file, such that values may not reach 100 million. Since PRMTOP uses coordinate array indices *N* = 3(*A −* 1), where *A* are the atom indices, *N* becomes too large beyond *∼*33 million atoms. NAMD’s existing PRMTOP parser rejects files whose values reach 10 million (seven digits), constraining atom counts to *∼*3.3 million.

Further, *tleap* pre-calculates coefficients for pairwise nonbonded interactions and stores them in the PRMTOP with limited precision. Since *tleap* does not have a parallel implementation, PRMTOP construction is unable to take advantage of modern multi-core computer architectures. Intractable wait times for systems comprising even several million atoms severely hampers researchers’ ability to debug, test, and effectively prepare models describing large systems. In contrast, NAMD natively runs CHARMMff and imports topology and force field details via a combination of X-PLOR format PSF files (produced by e.g., *psfgen*) and parameter files in either X-PLOR or CHARMM format. Notably, X-PLOR format PSF files use atom type names rather than numeric type indices. For atom counts beyond 10 million where memory optimization becomes important, NAMD can alternatively utilize the JS binary file format instead of the text based PSF; *psfgen* can also prepare JS files.

To fully merge the capabilities of AMBERff and NAMD, enabling biomolecular simulations of up to two billion atoms, we refactor AMBERff to produce CHARMM format force field files (i.e., TOP, RTF, PRM, STR) that interface seamlessly with *psfgen* and NAMD. Our approach allows the straightforward production of X-PLOR format PSF and JS files that faithfully reproduce potential energies and system properties calculated using parameters read from PRMTOP. The following sections describe the modifications required to represent AMBERff in CHARMM format for import into *psfgen* and ultimately NAMD. Table 1 lists relevant AMBERff file types and their NAMD analog for the described implementation.

**Table 1:** AMBERff file types/builders and their NAMD analog for this implementation.

|  | AMBER | NAMD |
| --- | --- | --- |
| Molecular topology, atom types, charges | LIB,OFF,PREP | TOP,RTF,STR |
| Parameters assigned via atom types | DAT,FRCMOD | PRM,STR |
| Typical software for system construction | *tLeap* | *psfgen* |
| Load topology, atom types, charges | LEAPRC in *tLeap* | TOP,RTF,STR in *psfgen* |
| Load parameter sets and modifications | LEAPRC in *tLeap* | NAMDRC in NAMD |
| Topology/parameters for simulation | PRMTOP | PSF,JS + NAMDRC |

### Force field equations

Within the framework of the present implementation, NAMD reads in AMBERff via a combination of PSF/JS and CHARMM format parameter files (i.e., in NAMD configuration: Amber off, ParaTypeCHARMM on). This means that AMBERff parameters are evaluated in the CHARMMff functional form. The AMBERff equation for potential energy is:

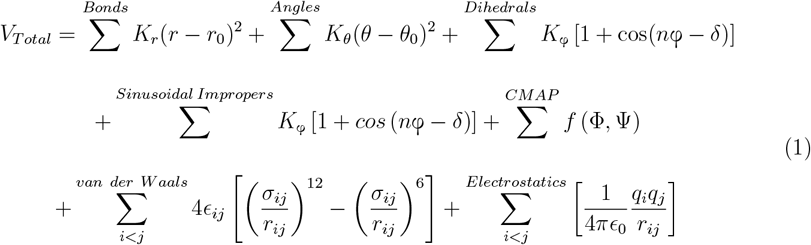

The CHARMMff equation for potential energy, as implemented in NAMD, is:

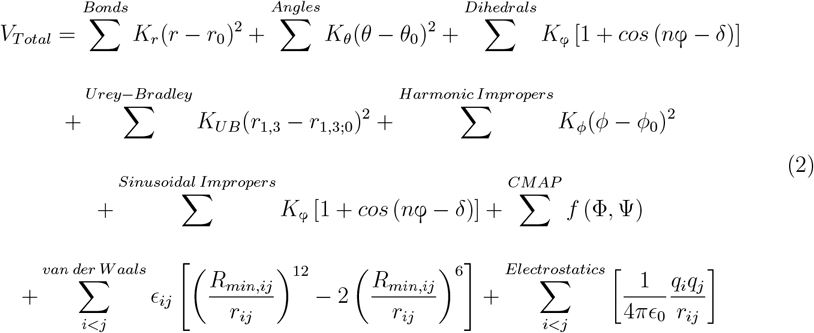

Given the mathematical similarity of the equations and the common units of parameters, AMBERff is readily substituted for CHARMMff in the NAMD integrator. In the absence of Urey-Bradley angle bending and harmonic improper dihedral angles, the contributions of these terms are eliminated, and Equation 2 reduces to Equation 1 when 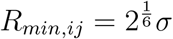. Sinusoidal improper dihedral angles receive the same treatment as proper torsions in AMBERff and are handled by the standard dihedral term.

### Atom types

Case sensitivity and the use of symbols, spaces, and numbers are key aspects of AMBERff atom type conventions. Although these details can be preserved in PSF and JS files, NAMD interprets CHARMM format atom types as capitalized, disregards symbols and spaces, and rejects leading numbers. In remedy, we introduce an atom type prefix system that encodes the information lost when ignoring case and symbols, which also conveniently handles numbers. Characters used to construct prefixes are summarized in Table 2. Prefixes comprise capital letters and contain as many characters as the original atom type. Each character of a prefix represents a corresponding character in the atom type it renames and denotes its nature: uppercase, lowercase, space, number, asterisk, plus sign, or minus sign. The prefix is followed by the alphanumeric portion of the original atom type, with any trailing symbols dropped. The new referenced atom types retain their uniqueness and are interpreted correctly by NAMD, allowing seamless import of AMBERff parameters.

**Table 2:**
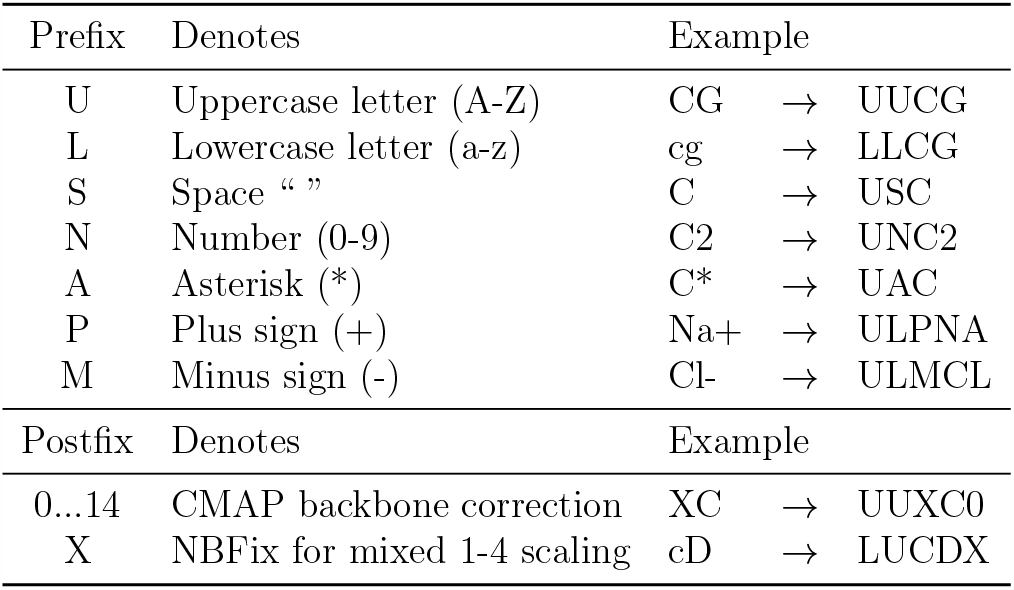
Prefixes and postfixes for AMBERff atom types in NAMD.

The most recent AMBERff for proteins, ff19SB,^22^ incorporates grid-based corrections to backbone torsions to more accurately reproduce *φ/ψ* conformational energy surfaces. ^23^ The CMAP term has identical functional forms in both AMBERff and CHARMMff (Eq. 1 and 2), such that it is readily handled by NAMD. However, AMBER format force field files encode CMAPs on a per-residue basis, while CHARMM format files encode them on the basis of dihedral type (i.e., combination of four atom types). To reconcile this difference, we introduce new atom types to distinguish the C*α* for each of the 16 unique CMAPs. The new atom types are simply duplicates with a numerical postfix appended, allowing CMAPs to be assigned to specific dihedrals without affecting usage of the original atom types in other contexts. For example, the alanine *φ* correction is applied to X*−*UUXC0*−*X*−*X, and atom type UUXC continues to be used in all other parameter declarations.

The most recent AMBERff for lipids, Lipid21,^24^ uses two different 1-4 nonbonded scaling factors (SCNB) to attenuate the van der Waals interactions between atoms separated by three bonds. All previous parameter sets use a single SCNB, although that value may be different across individual force fields. To support the use of multiple scaling factors within the same system, colloquially referred to as mixed scaling, AMBER format files assign SCNB on a per-dihedral basis. These values are written in the PRMTOP and applied as appropriate by the engine. However, CHARMM format force field files assign SCNB on the basis of atom type, providing pre-scaled Lennard-Jones parameters in the nonbonded entries. The latter approach can break down in a mixed scaling scenario if some atoms participate in multiple 1-4 interactions and require different SCNB for each. To address these cases, we introduce new types to distinguish atoms whose nonbonded scaling depends on interaction partner. We use CHARMM’s NBFix to correct the Lennard-Jones parameters for 1-4 pairings involving these atoms. The new atom types are simply duplicates with an X postfix appended, and original atom types remain unaffected.

### Dihedral declarations

The philosophy of the present implementation is to accurately reproduce the potential energies and ultimately the biophysical properties predicted by AMBERff in its native engine. To achieve full compatibility and energy agreement, it was occasionally necessary to alter parameter declarations, although not parameter values. These modifications account for data manipulation by *tleap* not emulated by *psfgen*. The result is that the described implementation ensures reproduction of AMBERff as encoded in the PRMTOP.

Notably, a number of dihedral declarations, particularly impropers, had to be revised. The definition of the improper *ϕ* angle is specified by the order of atoms in the declaration. There are a variety of circumstances that trigger *tleap* to reorder the atoms (e.g., A*−*B*−*C*−*D becomes D*−*A*−*C*−*B), thus redefining *ϕ*. Because *psfgen* does not make an equivalent change, it uses the former definition, while *tleap* uses the latter, leading to discrepancies in energies calculated with PRMTOP versus PSF. To address these issues, the described implementation includes dihedral declarations that correspond to those assigned by *tleap* for all circumstances. Amendments were made on a case-by-case basis either by revising atom order, adding duplicate declarations with the alternative atom order, or by introducing patches to correct atom order for special-case scenarios, as described below.

Also, following from AMBERff’s liberal use of wildcards in dihedral declarations, in rare circumstances, some parameters were found to be accidentally misallocated or overwritten by *tleap*. For such cases, it was necessary to modify parameter values to match, not those found in the AMBERff files, but those actually assigned by *tleap*. All such modifications are clearly documented within each respective force field file distributed with the implementation.

### Residue names and patches

In the present implementation, all AMBERff atom and residue names have been preserved. With the exception of histidine, both AMBERff and CHARMMff use the three-letter residue naming scheme formalized by the RCSB Protein Data Bank (PDB) ^25,26^ for standard amino acids. Consistent with AMBERff, histidine is here labeled HIE, HID, or HIP to denote the presence of hydrogen at the *E, δ*, or both *E* and *δ* positions. Other alternative, non-terminal protonation states are likewise handled with residue names, such as ASH/GLH for neutral aspartic/glutamic acid and LYN for neutral lysine.

While non-standard residue names included in AMBERff are read correctly by *psfgen* from the input PDB file, the implementation also includes patches to introduce them based on modification of their corresponding default residues, as is done in CHARMMff. Patches are also included for disulfide linkages (i.e., DISU introduces CYX pairs) and all available terminal capping residues. For the AMBER-DYES^27,28^ fluorophore force field, patches were introduced for bonding all possible linker-dye combinations. Additional patches were introduced to handle special cases where conditional reordering of improper dihedral declarations is necessary to produce correct potential energies. Such cases include PRO-PRO linkages in ff15ipq,^29,30^ certain C-terminal combinations in ff19SB, ^22^ and the first adenine residue of any sequence in OL3, OL15, and BSC1.^31–33^ All special-case patches and situations requiring them are described in the implementation documentation for the respective force field.

### Resource files

Many AMBER force fields are based on augmentation or modification of previous parameter sets, as with ff99, ^34^ superseded by ff99SB, ^35^ followed by ff99SB-ildn. ^36^ By AMBER convention, original parameter sets are stored as DAT files and modifications as FRCMOD files, with resource files (i.e., LEAPRC) provided to load all relevant components for PRMTOP construction with *tleap*. While topology information is encoded in PSF and JS files by *psfgen*, NAMD loads parameter details at runtime via X-PLOR or CHARMM format files. To preserve AMBER convention, original parameter sets and their modifications are provided as separate files, and a NAMDRC resource file is introduced to facilitate import of the relevant components of a given force field in the appropriate order.

### Usage

Usage of this implementation of AMBERff in *psfgen* and NAMD is analogous to that of CHARMMff. As indicated in Table 1, relevant TOP, RTF, and STR files for the desired force field(s) are loaded into VMD’s^20^ *psfgen* plugin for system construction. Segments are built and patches applied according to normal *psfgen* operation. The system may be immersed in a solvent box and ions added, either for neutralization or to effect a salt concentration, using newly AMBER-cognizant *solvate* and *autoionize* plugins. Consistent with the CHARMMff approach, solvent and ion parameters are presented in STR files, which contain both molecular topology and parameter information. Table 3 lists the force fields available for use with NAMD via this implementation at the time of publication.

**Table 3:**
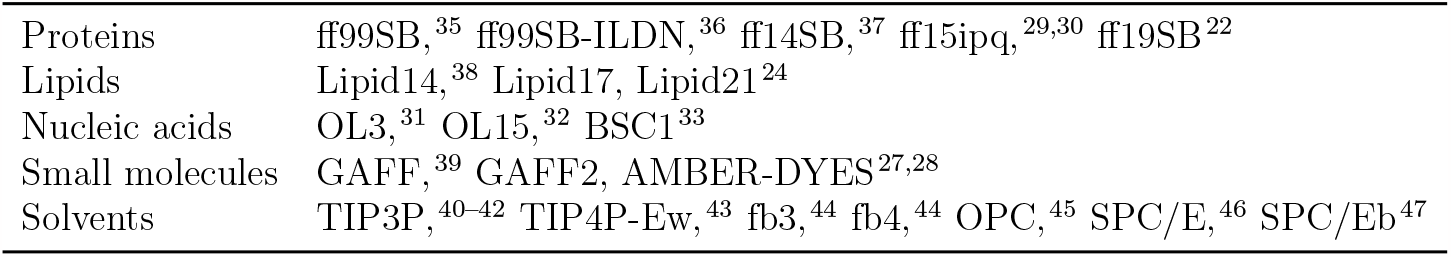
AMBERff for NAMD available at time of publication.

|  |  |
| --- | --- |
| Proteins | ff99SB,<sup>35</sup> ff99SB-ILDN,<sup>36</sup> ff14SB,<sup>37</sup> ff15ipq,<sup>29,30</sup> ff19SB<sup>22</sup> |
| Lipids | Lipid14,<sup>38</sup> Lipid17, Lipid21<sup>24</sup> |
| Nucleic acids | OL3,<sup>31</sup> OL15,<sup>32</sup> BSC1<sup>33</sup> |
| Small molecules | GAFF,<sup>39</sup> GAFF2, AMBER-DYES<sup>27,28</sup> |
| Solvents | TIP3P,<sup>40-42</sup> TIP4P-Ew,<sup>43</sup> fb3,<sup>44</sup> fb4,<sup>44</sup> OPC,<sup>45</sup> SPC/E,<sup>46</sup> SPC/Eb<sup>47</sup> |

An X-PLOR format PSF or JS file is written and provided to NAMD, along with initial coordinates, and the newly-introduced NAMDRC file is sourced. The NAMDRC, analogous to AMBERff’s LEAPRC, loads relevant PRM and STR parameter information for a given force field, as well as sets the simulation configuration: Amber off, ParaTypeCHARMM on. The value of 1-4scaling should be set to 1*/*1.2 = 0.833333 to assign the correct 1-4 electrostatic scaling factor for AMBERff proteins, lipids, and nucleic acids. Example *psfgen* and NAMD configuration scripts for the MD simulations discussed in this work are provided in Supplemental Information. These examples include additional configuration details required to preserve the integrity of AMBERff when using NAMD.

## Results

### Validation of potential energies

To demonstrate the validity of the described AMBERff implementation in NAMD, results were compared for three scenarios: (i) PRMTOP in AMBER22^48^ *pmemd*.*cuda*, (iii) XLPOR format PSF with CHARMM format parameter files in NAMD 2.14, and (iii) the latter in NAMD 2.14 compiled with *pmemd*.*cuda*’s Coulomb’s constant, referred to as NAMBER. Configuration settings were selected to minimize algorithmic differences between the engines. ^49,50^ Large cutoffs were employed to force direct-space calculation of electrostatic interactions, as well as prevent the truncation or long-range correction of van der Waals interactions. The AMBER and NAMD configuration scripts utilized in testing are provided in Supplemental Information.

For single-point potential energy validation, protein test systems included ensembles of all possible tripeptide sequences available in each force field, considering all possible terminal capping residues. Lipid test systems included ensembles of all head groups with every possible tail in the sn1 and sn2 positions, covering all lipid molecules available in each force field. Steroids treated as complete residues (e.g., cholesterol) were also included in the lipid ensembles. Nucleic acid test systems included ensembles of all possible trinucleotide DNA/RNA sequences available in each force field, constructed with AMBER’s nucleic acid builder (NAB).^51^ In the case of AMBER-DYES, test systems included all individual residues, as well as all possible linker-fluorophore combinations.

Figure 1 shows the average and maximum single-point potential energy deviations comparing results from PRMTOP/*pmemd*.*cuda* with PSF/NAMD (green) and PSF/NAMBER (blue).

**Figure 1:**
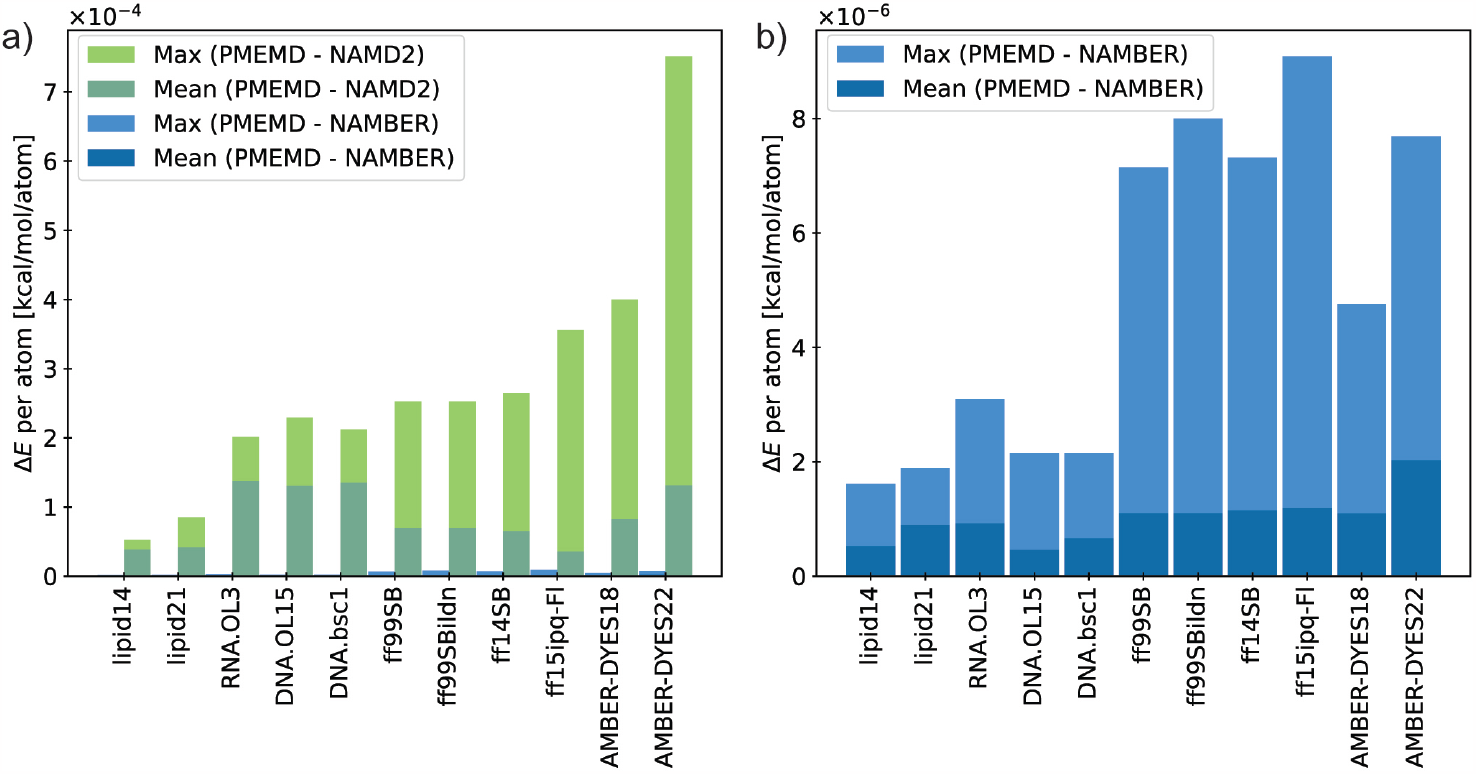
Comparing potential energies calculated with the AMBER engine versus NAMD (green) and NAMBER (blue) for various force fields. **a)** Per-atom single-point energy deviations. **b)** Deviations decrease by two orders of magnitude using NAMBER (NAMD compiled with AMBER’s Coulomb’s constant). Owing to an update in the AMBER-DYES fluorophore parameters^52,53^ between the release of AMBER18 and AMBER22, both versions are included and referred to as AMBER-DYES18 and AMBER-DYES22, respectively.

The average deviations for all refactored force fields are less than 1.5 *×* 10^*−*4^ kcal*/*mol per atom for running AMBERff in NAMD versus its native engine (Fig. 1a). Small differences arise from the angle terms, due to the precision of internal constants used to convert between degrees and radians in the respective engines. Consistent with previous reports comparing MD simulation software,^54^ disparities in calculated energies are primarily attributable to the electrostatics terms.

AMBER22 *pmemd*.*cuda* and NAMD 2.14 use Coulomb’s constant 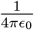 equal to 332.0522173 and 332.0636 kcal/mol Å *e*^*−*2^, respectively. ^49^ Further, the AMBER engine reads the values of atomic charges multiplied by 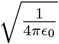 from the PRMTOP, while NAMD calculates these values at runtime and stores them in memory with higher precision. Using NAMD compiled with *pmemd*.*cuda*’s Coulomb’s constant decreased the average single-point potential energy deviations by two orders of magnitude, to less than 2.1 *×* 10^*−*6^ kcal*/*mol per atom (Fig. 1b). These modest differences are not expected to affect thermodynamics computations to within the statistical error generally accepted by simulation studies. ^54^

### Validation of biophysical properties

To provide confidence that the deviations quantified above do not significantly alter predicted biophysical properties, test cases for protein, lipid and nucleic acid systems were evaluated. For each system, a PRMTOP was prepared using *tleap* from AMBER22^48^ and an X-PLOR format PSF was prepared using *psfgen* from VMD 1.9.3. ^20^ Simulations were performed using *pmemd*.*cuda* and the distributed version of NAMD 2.14. Configuration settings were selected to minimize algorithmic differences between the engines.^49,50^ The configuration scripts describing simulation settings are provided in Supplemental Information. Simulations were run in triplicate for both engines. All trajectory analysis was carried out using VMD.

Minimization utilized the hybrid conjugate gradient/steepest descent algorithm, and dynamics were propagated in the isothermal-isobaric ensemble (NPT). Velocities were initialized using a different random seed in each engine. Bonds containing hydrogen atoms were constrained, facilitating a timestep of 2 fs. Temperature was controlled using the Langevin thermostat with a friction coefficient of 1 ps^*−*1^. Pressure was controlled at 1 bar using the Berendsen barostat with isotropic scaling and a relaxation time of 1 ps. Long-range electrostatics were calculated using particle-mesh Ewald (PME) with cubic interpolation and a direct space tolerance of 1*×*10^*−*6^. Nonbonded interactions were cutoff at 8 Å with no switching function, applying an analytical correction to approximate the long-range Lennard-Jones potential. Trajectory frames were saved every 5,000 steps.

For robust statistical comparison of biophysical property distributions obtained from simulations, we applied the Jensen-Shannon Divergence (JSD) test. The bounded value 0 *≤ JSD ≤* 1 serves as a measure of similarity, with zero signifying that two distributions are identical and one signifying that they are maximally different. The JSD between probability distributions *P* and *Q* is defined as:

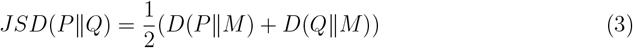

where 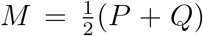 is the average distribution and *D* represents the Kullback-Leibner Divergence (KLD), defined as:

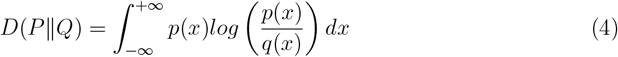

### Test case: Ubiquitin

Ubiquitin was examined as a test case for protein force fields (Fig. 2a, inset). The highresolution crystal structure (PDB ID: 1UBQ^55^) was assigned hydrogen atoms corresponding to pH 7.0 using the PROPKA^56,57^ method in PDB2PQR,^58,59^ while maintaining crystallographic waters. The system was immersed in a 76*×*76*×*76 Å^3^ box of TIP3P^40^ solvent containing 150 mM NaCl, ^60^ and the ff14SB force field^37^ was applied. The system was subjected to energy minimization for 5,000 steps and heated from 50 K to 310 K over 5 ns, while maintaining backbone restraints. Restraints were gradually released over 5 ns, and the systems were equilibrated for 150 ns. Production simulations were run in triplicate for 500 ns for a total of 1.5 μs cumulative sampling. For analysis, trajectories were aligned to the crystal structure based on the C*α* trace.

**Figure 2:**
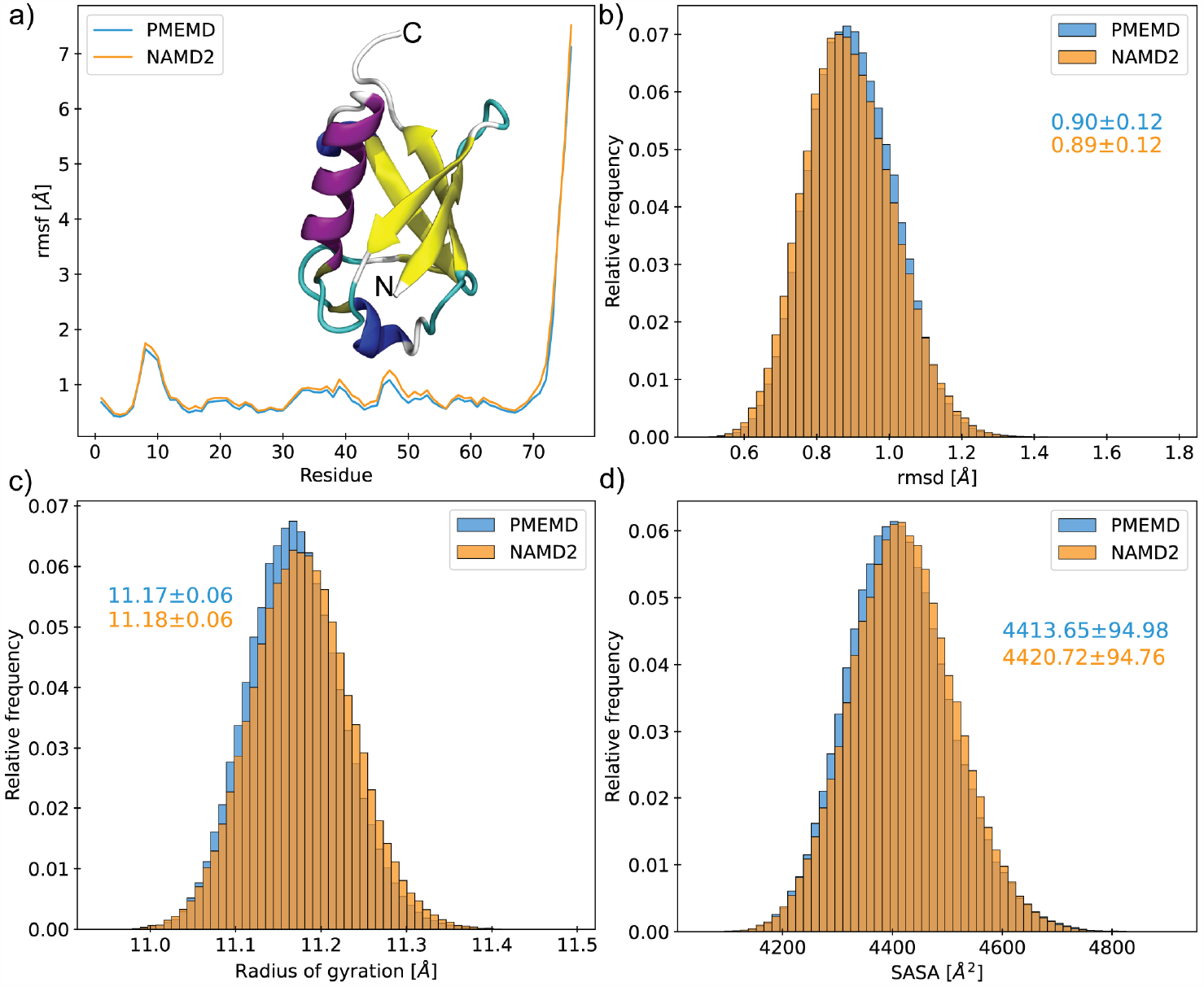
Assessing conservation of biophysical properties between simulations of ubiquitin (inset a) performed in AMBER (blue) and NAMD (orange) using the ff14SB force field. **a)** C*α* root-mean-square fluctuation. Frequency distributions for **b)** backbone root-mean-square deviation, **c)** radius of gyration, and **d)** solvent accessible surface area.

Comparable per-residue root-mean-square fluctuation (RMSF) profiles were produced by ubiquitin simulations with AMBER and NAMD (Fig. 2a), indicating similar biophysical behavior regardless of engine. Distributions of backbone root-mean-square deviation (RMSD), radius of gyration (R*g*), and solvent accessible surface area (SASA), all excluding highly flexible C-terminal residues 72-76, are shown in Figure 2b-d. These distributions display substantial overlap, with averages differing by *<*0.15*σ*. The magnitude of *JSD* for each property is on the order of 10^*−*3^ (Table S1), with values close to zero indicating very high, statistically significant similarity. The *JSD* between replicate simulations performed with either AMBER or NAMD is on the order of 10^*−*2^ (Fig. S1), such that inter-engine divergence is minimal and likely the result of limited sampling.

### Test Case: Dickerson-Drew dodecamer

The B-DNA sequence CGCGAATTCGCG, commonly referred to as the Dickerson-Drew Dodecamer (DDD),^61^ was examined as a test case for nucleic acid force fields (Fig. 3a). A model of DDD was constructed using AMBER’s nucleic acid builder (NAB).^51^ The system was immersed in a 76*×*76*×*76 Å^3^ box of TIP3P^40^ solvent containing 150 mM NaCl, ^60^ and the OL15 force field^32^ was applied. The system was subjected to energy minimization for 500 steps and heated from 50 K to 310 K over 5 ns, while maintaining backbone restraints. Restraints were gradually released over 5 ns. Production simulations were run in triplicate for 500 ns for a total of 1.5 μs cumulative sampling. For analysis, the *do x3DNA*^62,63^ plugin of VMD was used to track 27 structural properties.

**Figure 3:**
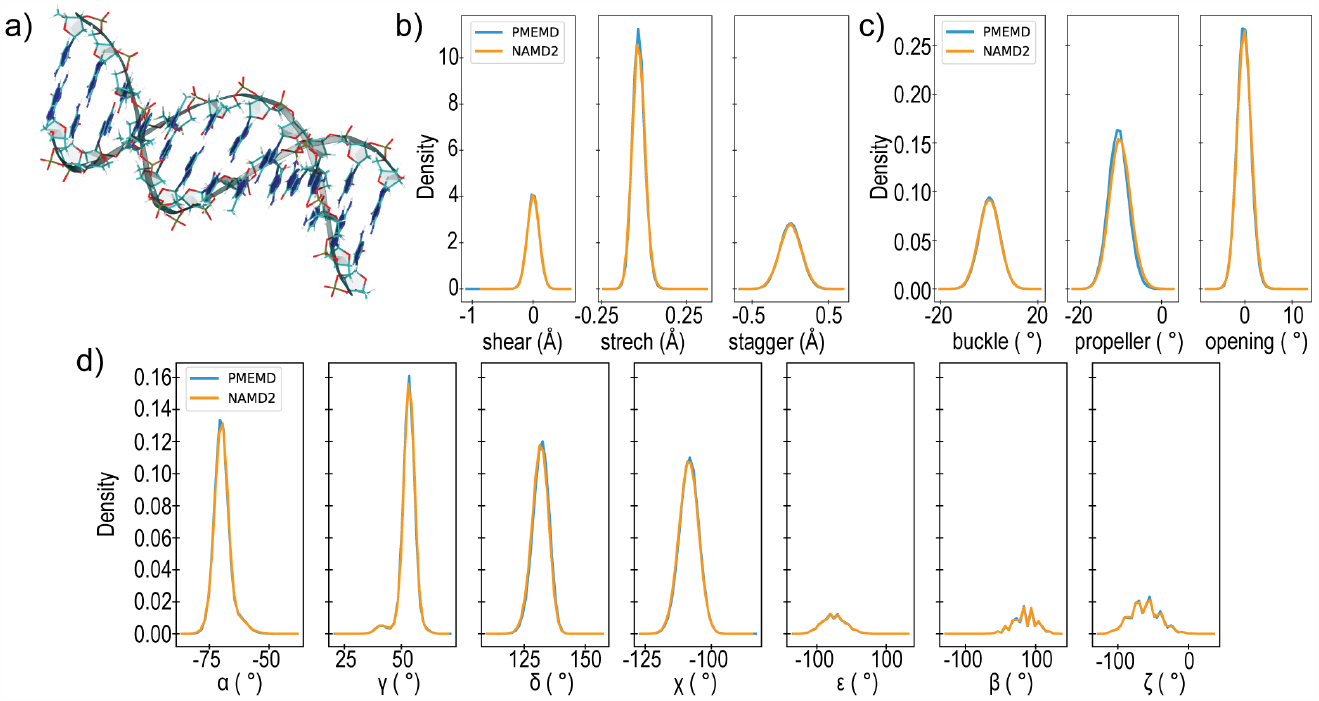
Assessing conservation of biophysical properties between simulations of **a)** the Dickerson-Drew dodecamer performed in AMBER (blue) and NAMD (orange) using the OL15 force field. Frequency distributions for **b)** radial base pair parameters, **c)** angular base pair parameters, and **d)** backbone dihedral angles.

Comparison of structural properties obtained from DDD simulations with AMBER and NAMD indicates similar biophysical behavior regardless of engine. Distributions of base pair parameters (Fig. 3b-c) and backbone dihedrals (Fig. 3d) display substantial overlap, and in most cases are indistinguishable. Terminal base pairs were excluded from analysis, as these are known to transiently unravel.^64^ A comprehensive presentation of the distribution data for all 27 structural properties, as well as their tabulated *JSD* values, are given in Supplemental Information. The magnitude of *JSD* for each structural property is on the order of 10^*−*3^ or less (Table S1), indicating very high, statistically significant similarity. The *JSD* between replicate simulations performed with either AMBER or NAMD is likewise on the order of 10^*−*3^ (Fig. S3). In some instances, for the timescale investigated, ensembles collected using different engines were more similar than those collected using the same engine.

### Test case: POPC membrane bilayer

A membrane bilayer composed of 1-palmitoyl-2-oleoyl-sn-glycero-3-phosphocholine (POPC) was examined as a test case for lipid force fields (Fig. 4a-b). A flat 76*×*76 Å^2^ bilayer patch was constructed using CHARMM-GUI. ^65^ TIP3P^40^ solvent containing 150 mM NaCl^60^ was added to produce a simulation box of 85 Å along the z-dimension, and the Lipid21 force field^24^ was applied. The system was subjected to energy minimization for 10,000 steps and heated from 60 K to 393 K over 1 ns. High-temperature dynamics at 393 K were performed over 10 ns to melt the lipid tails. The system was then cooled over 1 ns to 303 K and then equilibrated for 100 ns. Three production simulations were forked from the equilibrated coordinates and each run for 500 ns for a total of 1.5 μs cumulative sampling.

**Figure 4:**
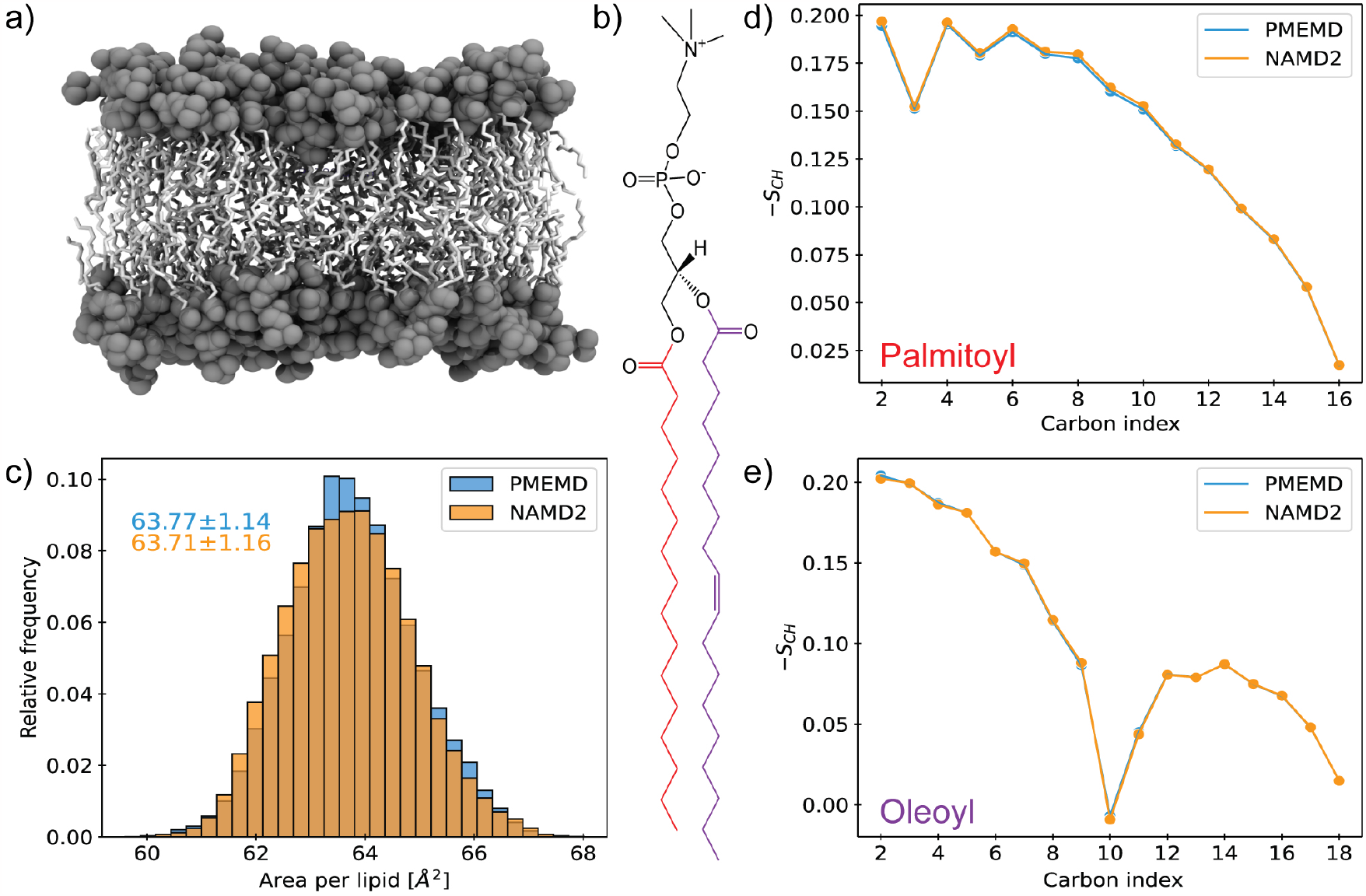
Assessing conservation of biophysical properties between simulations of **a)** a membrane bilayer composed of **b)** POPC performed in AMBER (blue) and NAMD (orange) using the Lipid21 force field. **c)** Frequency distribution for area per lipid. Order parameters (S_CH_) for the **d)** palmitoyl (red) and **e)** oleoyl (purple) tails.

Area per lipid (APL) distributions (Fig. 4c) display substantial overlap for POPC simulations in both AMBER and NAMD. The APL value of 63.7 Å^2^ obtained here matches well with independent simulations performed at 303 K (63.9 Å^2^),^24^ as well as reported experimental measurements (64.3 Å^2^).^66^ The magnitude of *JSD* for APL distributions is on the order of 10^*−*3^ (Table S1), indicating very high, statistically significant similarity. The *JSD* between replicate simulations performed with either AMBER or NAMD is on the order of 10^*−*2^ (Fig. S4), demonstrating again that the choice of engine does not meaningfully impact calculation results, particularly for common sampling timescales. In addition, the lipid tail order parameters for both paltmitoyl (Fig. 4d) and oleoyl (Fig. 4e) closely correspond, with the largest deviation being *<*2.5 *×* 10^*−*3^. Altogether, these results indicate similar biophysical behavior of the bilayer, regardless of engine.

### Solvent models

Generally, force fields are validated with and/or shown to more closely reproduce experimental properties with specific solvent models. Further, solvent and ion parameters are intrinsically linked. Unlike in CHARMMff, where force fields are inherently tied to solvent models owing to the charge parameterization scheme, solvent models are generally interchangeable in AMBERff. Common solvent models were validated for the described implementation by calculating average bulk properties obtained from simulations in AMBER and NAMD. Systems consisted of 27*×*27*×*27 Å^3^ pure solvent boxes constructed using *tleap* for the AMBER simulations and *psfgen* using the same coordinates for NAMD simulations. Systems were subjected to energy minimization for 1,000 steps and equilibrated for 100 ps. Thirty 100 ps simulation replicates were forked from the equilibrated coordinates, exploring either the isothermal-isobaric (NPT) or microcanonical (NVE) ensembles. NPT simulations were used to evaluate solvent density and dielectric constant *E*, and NVE simulations were used to assess self-diffusion coefficient *D*.^45,67^ Average values calculated over simulation replicates for these quantities were found to match closely for all solvent models tested (Fig. 5a-c), indicating appropriate bulk behavior.

**Figure 5:**
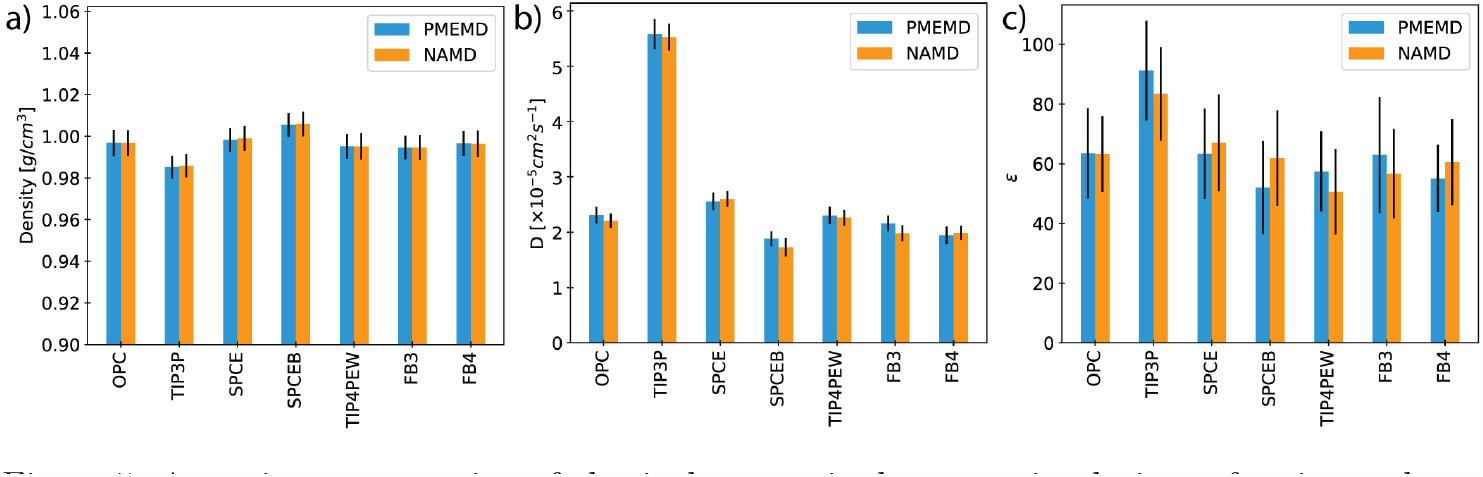
Assessing conservation of physical properties between simulations of various solvent models in AMBER (blue) and NAMD (orange). Calculated values for solvent **a)** density, **b)** self-diffusion coefficient, and **c)** dielectric constant. Error bars represent standard deviation.

## Discussion

Recently, AMBERff has been applied for the first time to simulate multimillion-atom systems in NAMD.^68,69^ These endeavors utilized either an early version of the described implementation, or extensive custom editing of the parameter/topology files to successfully load the force field into the engine. In the latter case, because input was encoded in PRMTOP rather than memory-optimized JS, simulation performance suffered despite access to a leadership-class supercomputer. ^69^ Memory-optimized NAMD is appropriate for systems larger than *∼*10 million atoms and essential for systems above *∼*30 million.^70^ The implementation presented here overcomes previous limitations to enable high-performance, massively parallel simulations with AMBERff encompassing up to two billion atoms, the current upper bound in NAMD2. Our results demonstrate that these simulations accurately conserve the biophysical behavior predicted by AMBERff in its native engine. The implementation produces equivalent results using NAMD2, GPU-accelerated NAMD2, and GPU-resident NAMD3.

During the past 15 years, numerous force field refinements have been spurred by validation testing over increasingly long simulation timescales afforded by novel enhanced sampling methods,^71,72^ GPU-acceleration^13,15^ and the special purpose ANTON machine.^73,74^ The application of AMBERff to biomolecular systems of increasing size and complexity will reveal new opportunities to advance classical force field development, expanding resolution of the computational microscope. Progressively larger and more detailed biomolecular structures emerging from the experimental sphere represent prime targets for high-impact discoveries driven by MD simulations. The rise of exascale computing and data analysis powered by machine learning positions researchers to examine the atomistic dynamics of massive biomolecular assemblies, already including intact viruses,^75^ small organelles, ^18^ and minimal cells. The implementation presented here broadens the computational technology available to investigate such viral and cellular machinery by allowing large-scale simulations to benefit from the decades of high-quality force field development that have culminated in AMBERff.

## Availability

The refactored force field files for AMBERff prepared and validated through this work, as well as AMBERff-cognizant *solvate* and *autoionize* plugins for VMD, are available from: https://github.com/Hadden-lab/AMBERff-in-NAMD.

## Acknowledgement

This work was inspired by early efforts of Professor Thomas E. Cheatham III, Professor Jeffery Klauda, and Dr. Xibing He to port AMBER force fields to the CHARMM and X-PLOR engines. This work was funded by University of Delaware Research Foundation and NSF award CBET-2232718. This work was supported by Delaware Advanced Research Workforce and Innovation Network (DARWIN), funded by NSF award OAC-1919839. Computer time on Anvil at Purdue University and Delta at National Center for Supercomputing Applications was provided by allocation BIO-230085 from Advanced Cyberinfrastructure Coordination Ecosystem: Services & Support (ACCESS) program. ACCESS is funded by NSF awards #2138259, #2138286, #2138307, #2137603, and #2138296. This work was supported by BioStore data storage resource made possible by NIH through Delaware IDeA Network of Biomedical Research Excellence, awards P20GM103446 and S10OD028725.

## Supplemental Information

**Table S1:** Jensen-Shannon Divergence values comparing biophysical property distributions.

| Property | $JSD(AMBER\vert \vert NAMD)$ |
| --- | --- |
| Protein test case: UBQ |  |
| RMSD | $1.86 \times 10^{-3}$ |
| RGYR | $4.89 \times 10^{-3}$ |
| SASA | $1.70 \times 10^{-3}$ |
| Nucleic acid test case: DDD |  |
| shear | $2.20 \times 10^{-4}$ |
| stretch | $4.35 \times 10^{-4}$ |
| stagger | $4.93 \times 10^{-4}$ |
| buckle | $3.03 \times 10^{-4}$ |
| propeller | $6.52 \times 10^{-3}$ |
| opening | $5.19 \times 10^{-4}$ |
| $\alpha$ | $6.75 \times 10^{-4}$ |
| $\beta$ | $1.25 \times 10^{-4}$ |
| $\gamma$ | $5.03 \times 10^{-4}$ |
| $\delta$ | $1.90 \times 10^{-3}$ |
| $\chi$ | $5.44 \times 10^{-4}$ |
| $\epsilon$ | $7.50 \times 10^{-5}$ |
| $\zeta$ | $7.50 \times 10^{-5}$ |
| shift | $2.45 \times 10^{-4}$ |
| slide | $1.29 \times 10^{-3}$ |
| rise | $5.22 \times 10^{-3}$ |
| tilt | $4.14 \times 10^{-4}$ |
| roll | $6.77 \times 10^{-4}$ |
| twist | $4.78 \times 10^{-3}$ |
| x-displacement | $3.47 \times 10^{-4}$ |
| y-displacement | $4.22 \times 10^{-4}$ |
| helical rise | $8.72 \times 10^{-4}$ |
| inclination | $9.52 \times 10^{-4}$ |
| tip | $4.48 \times 10^{-4}$ |
| helical twist | $3.17 \times 10^{-3}$ |
| major groove | $7.46 \times 10^{-3}$ |
| minor groove | $2.21 \times 10^{-3}$ |
| Lipid test case: POPC |  |
| APL | $2.14 \times 10^{-3}$ |

**Figure S1:**
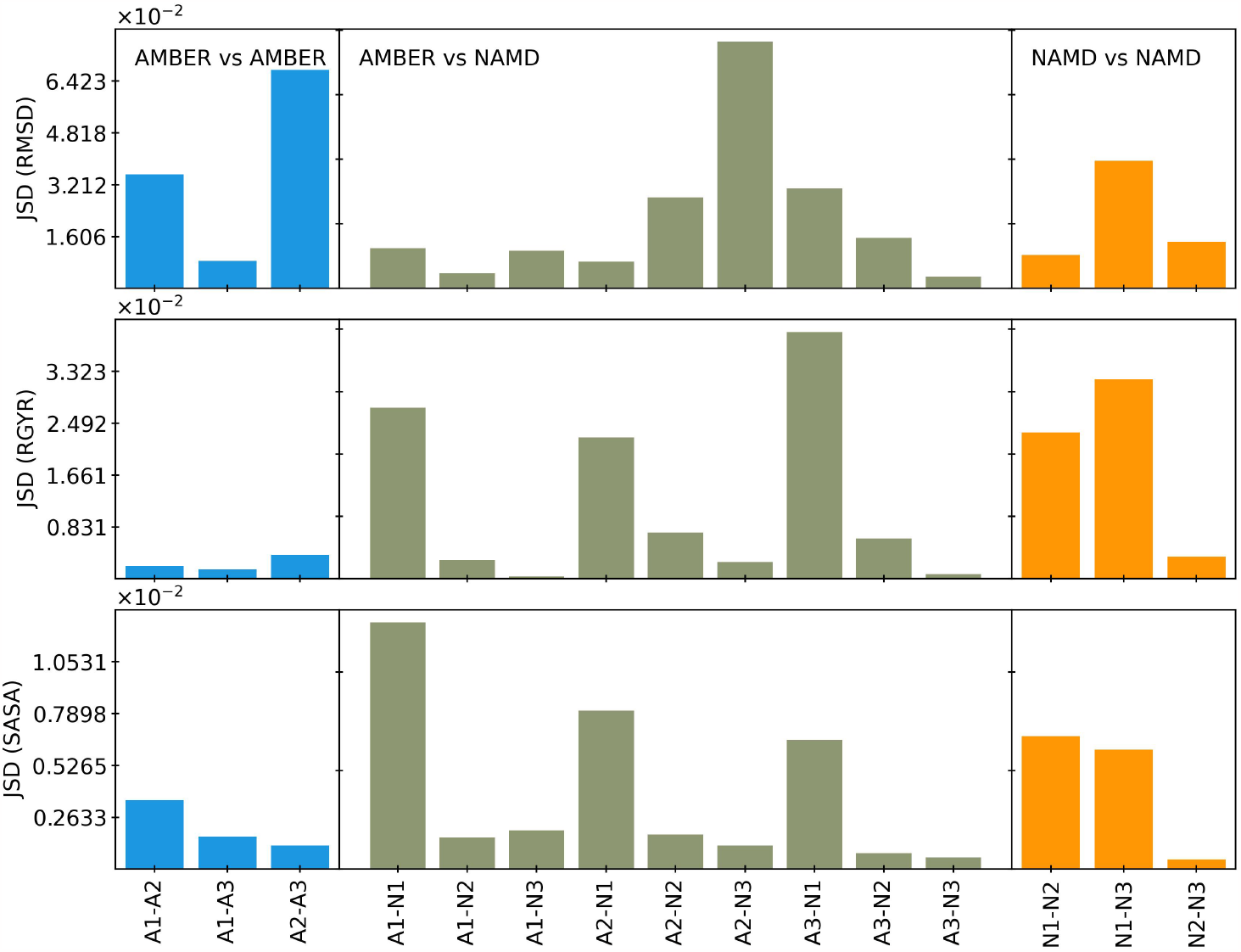
Jensen-Shannon Divergence values comparing biophysical property distributions for individual protein simulations.

**Figure S2:**
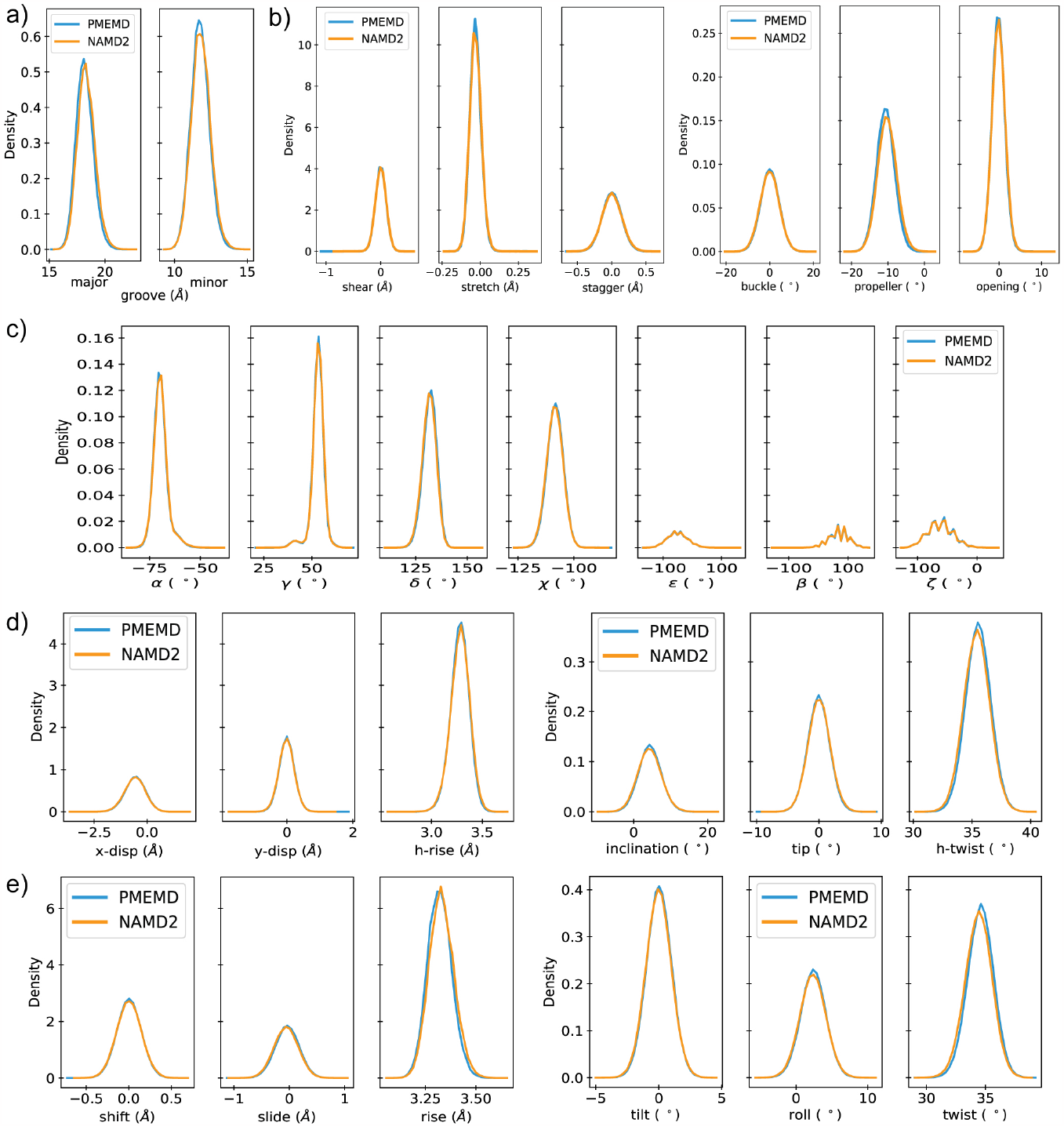
Distributions obtained from simulations in AMBER (blue) and NAMD (orange) for the complete set of 27 structural parameters tracked for the Dickerson-Drew dodecamer. Structural parameters include: **a)** major and minor groove widths, **b)** base pair parameters (shear, stretch, stagger, buckle, propeller and opening), **c)** backbone dihedral angles, **d)** helical base step parameters (displacement along x-axis and y-axis, helical rise, inclination, tip, and helical twist), and **e)** base step parameters (shift, slide, rise, tilt, roll, and twist).

**Figure S3:**
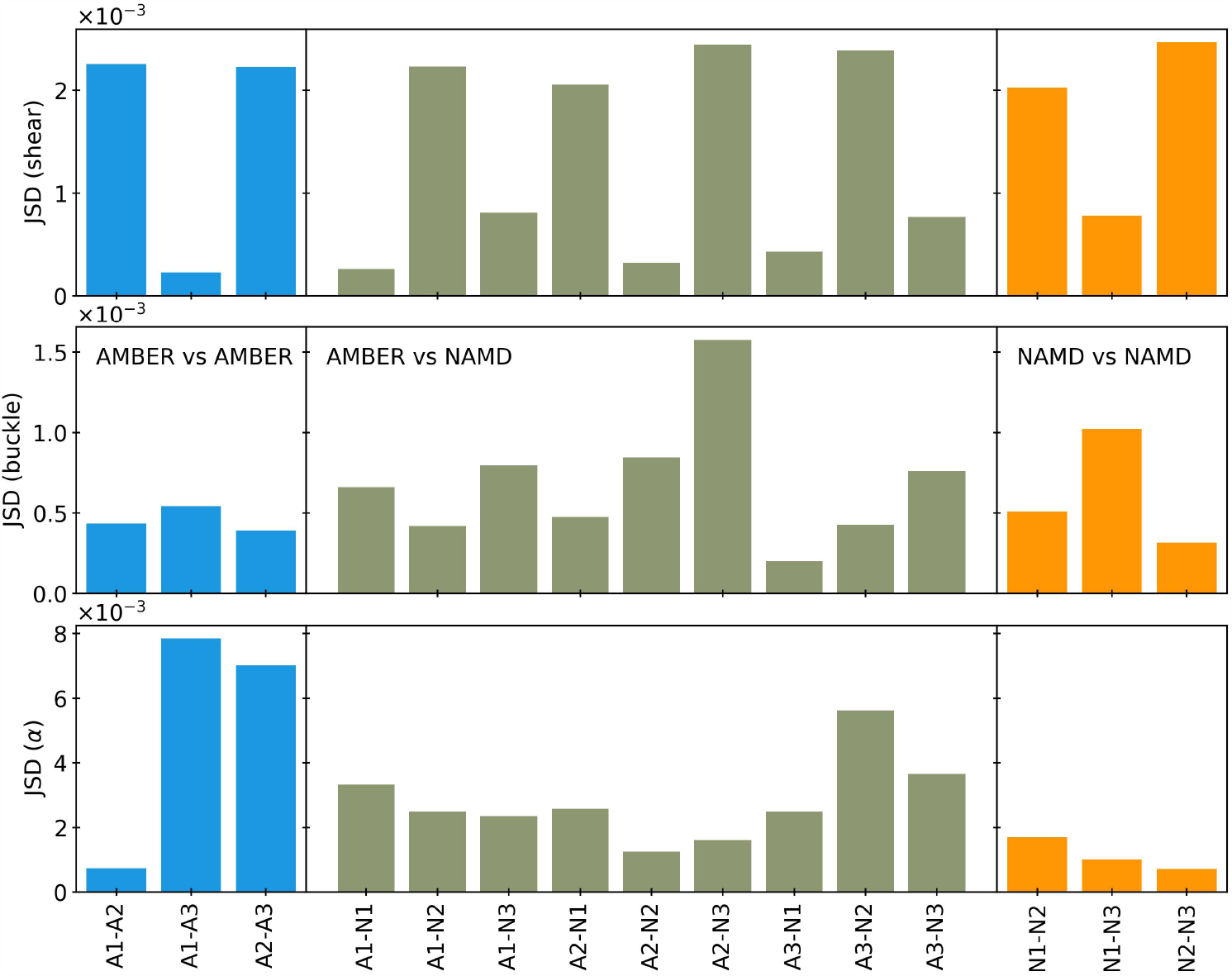
Jensen-Shannon Divergence values comparing biophysical property distributions for individual nucleic acid simulations.

**Figure S4:**
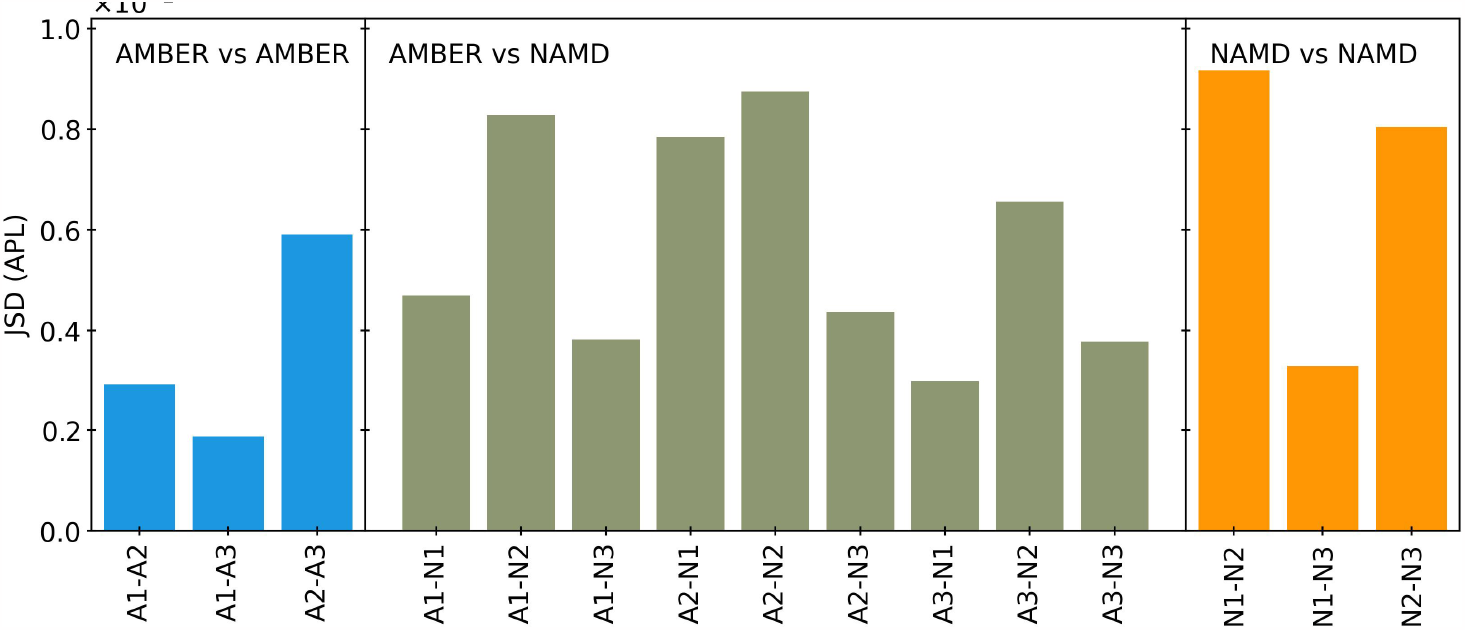
Jensen-Shannon Divergence values comparing biophysical property distributions for individual lipid simulations.

## Scripts

### Example VMD script using *psfgen*

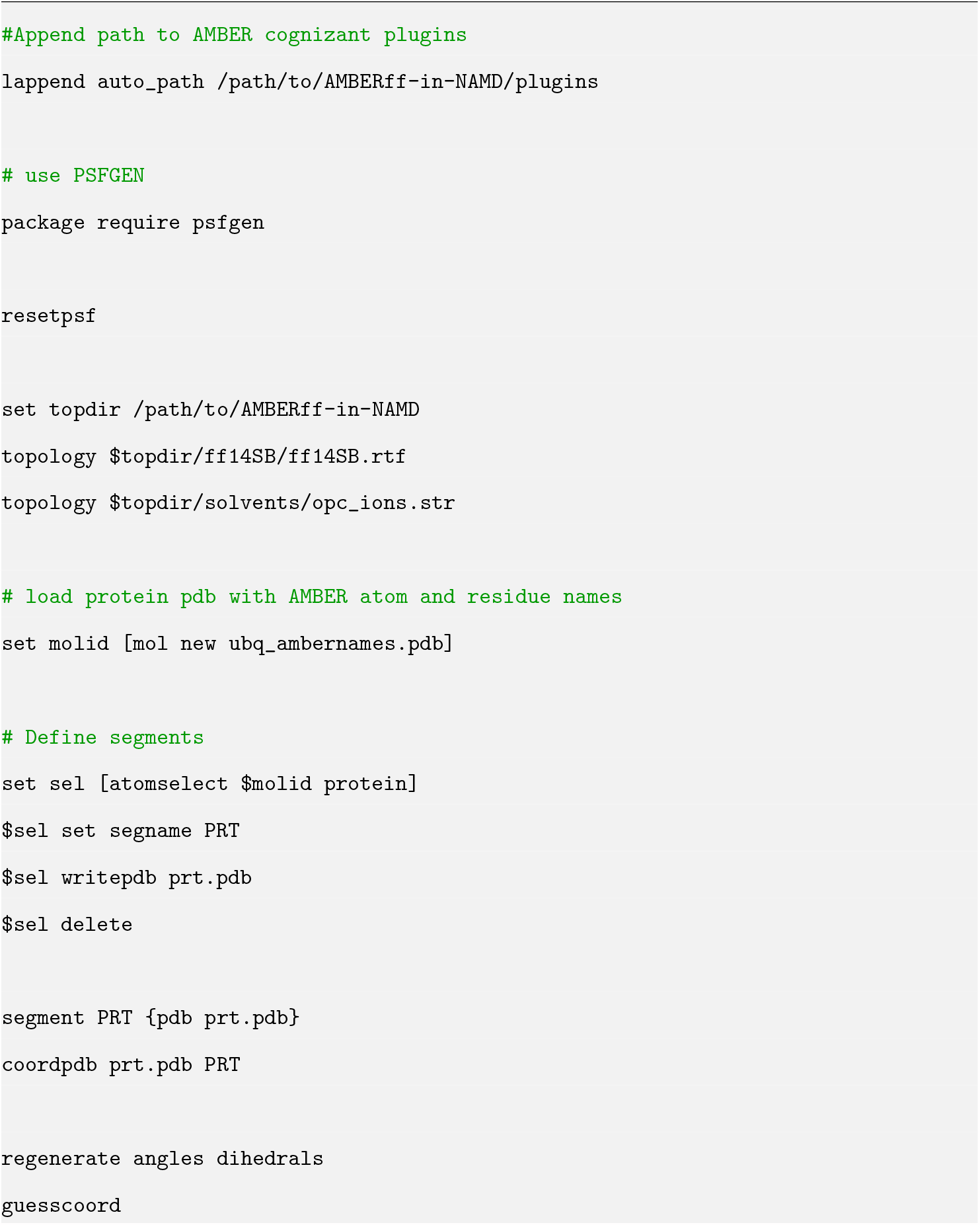
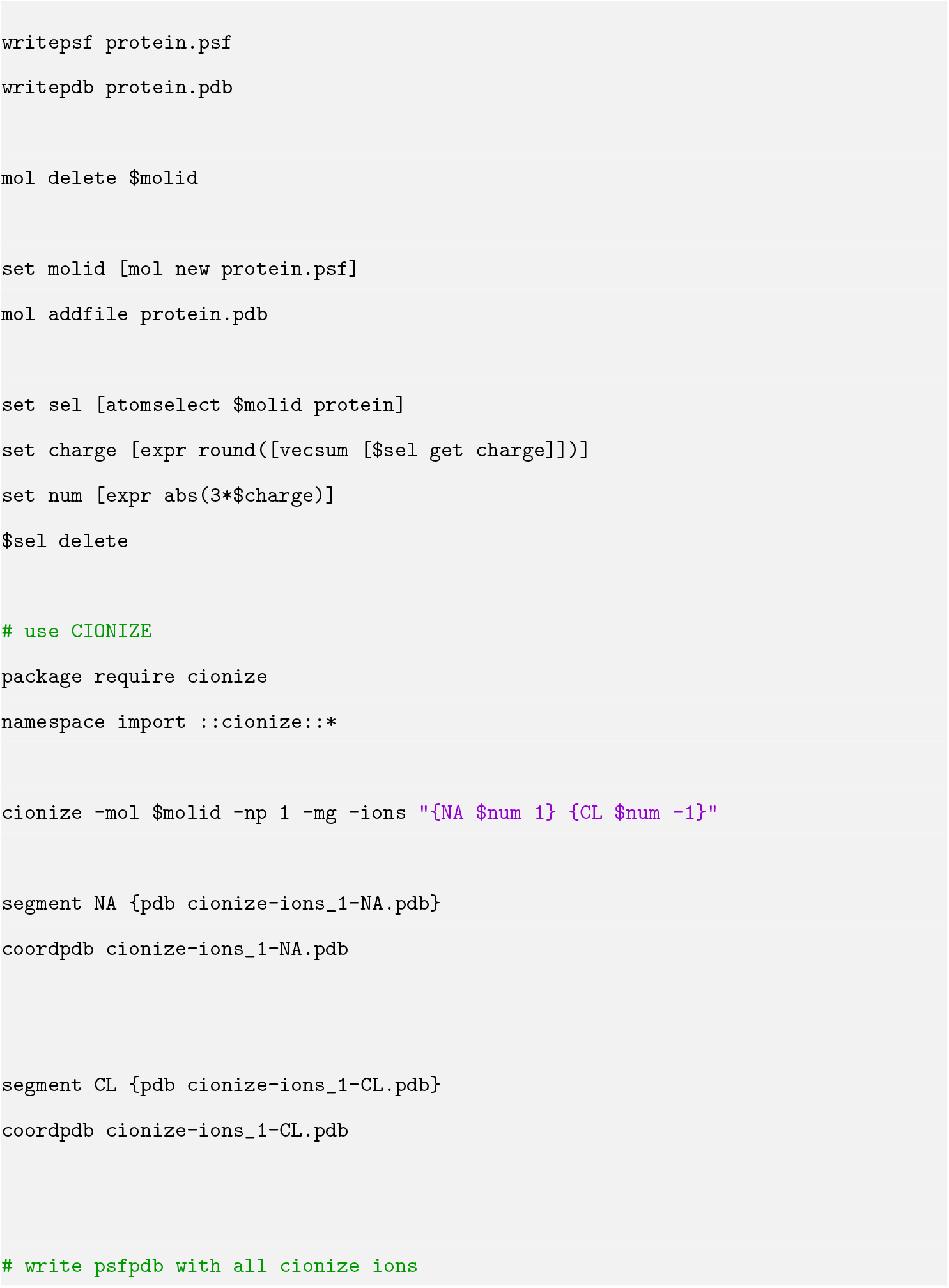
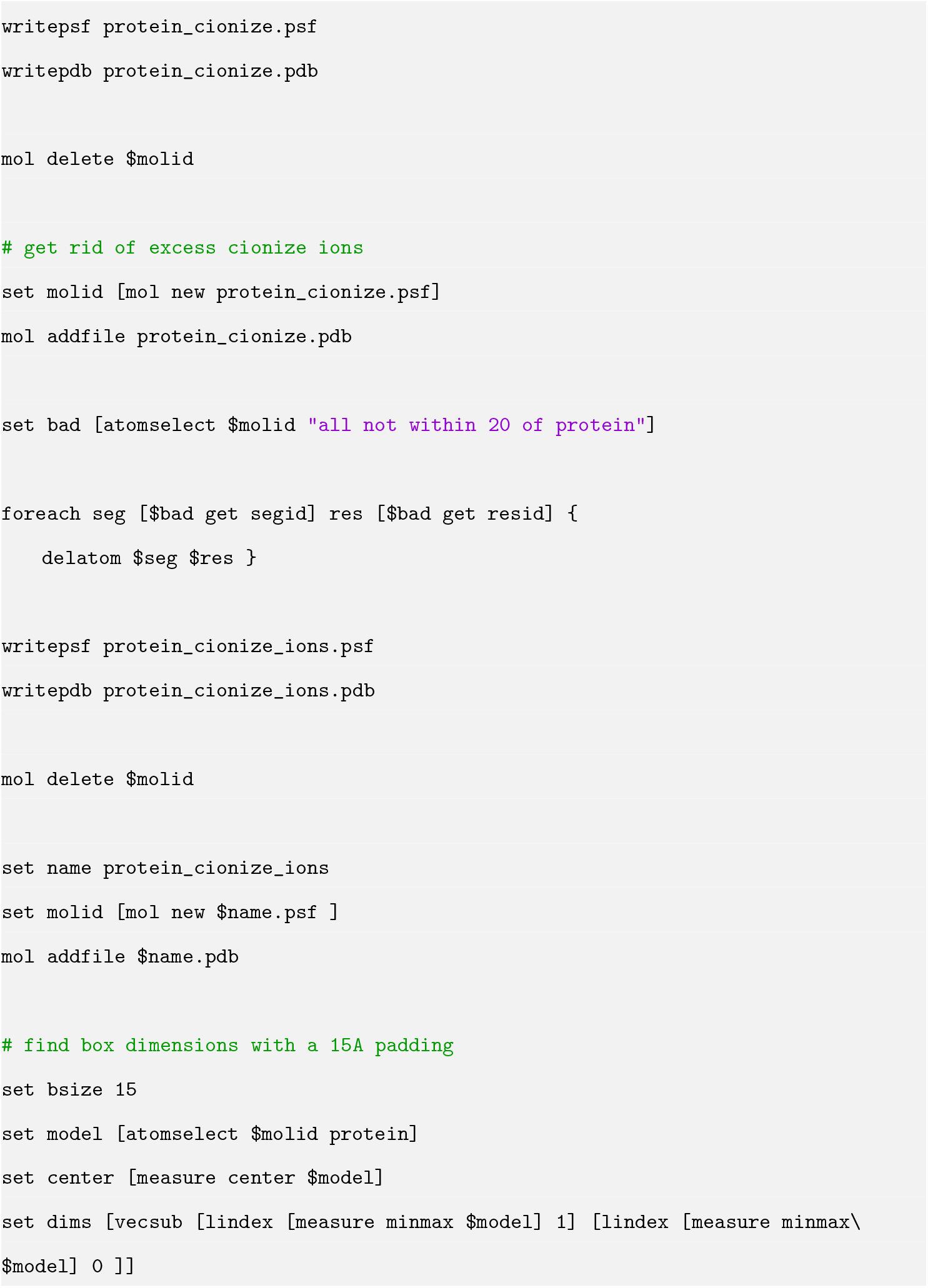
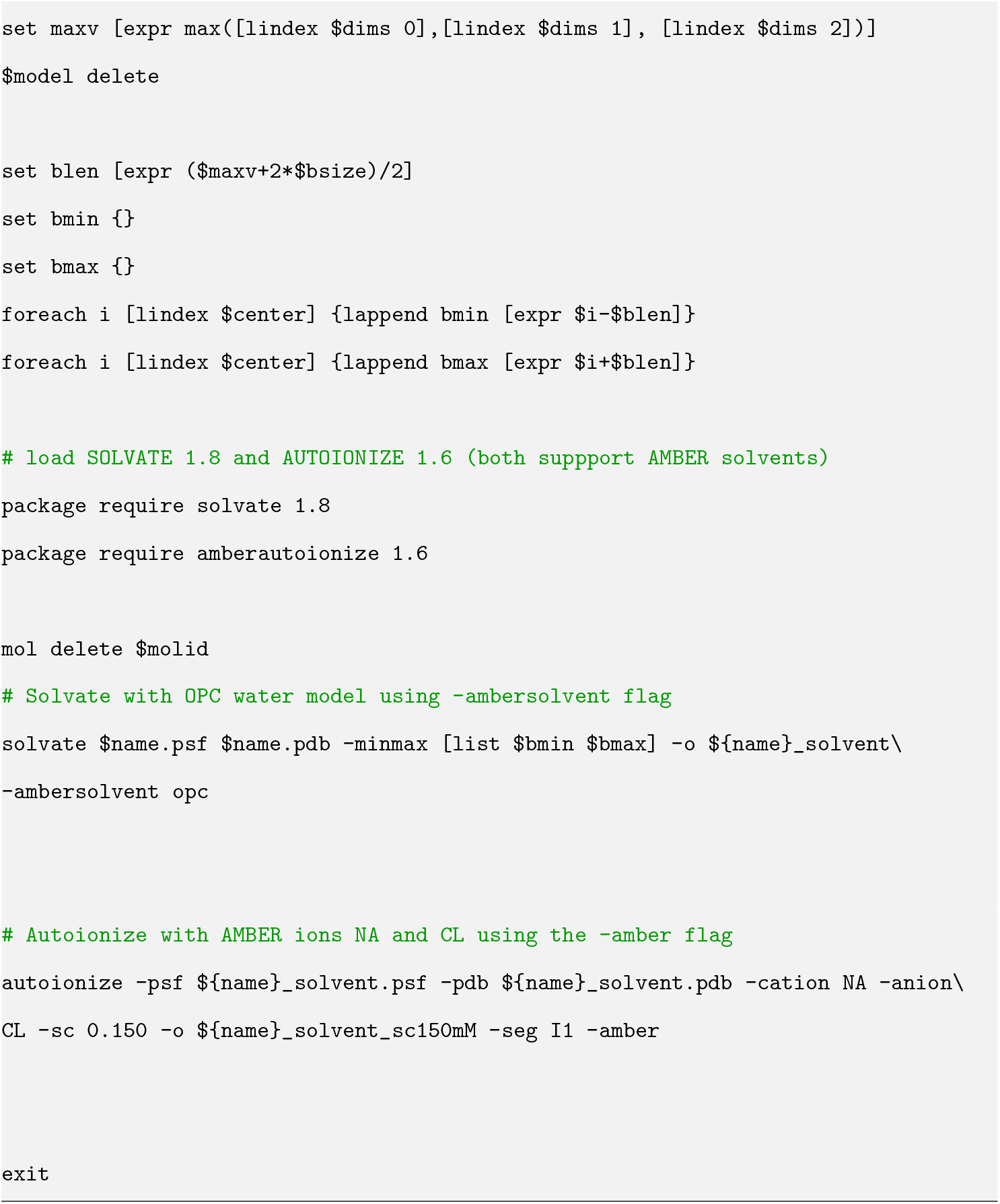

### PMRTOP in AMBER: Single point energy configuration file

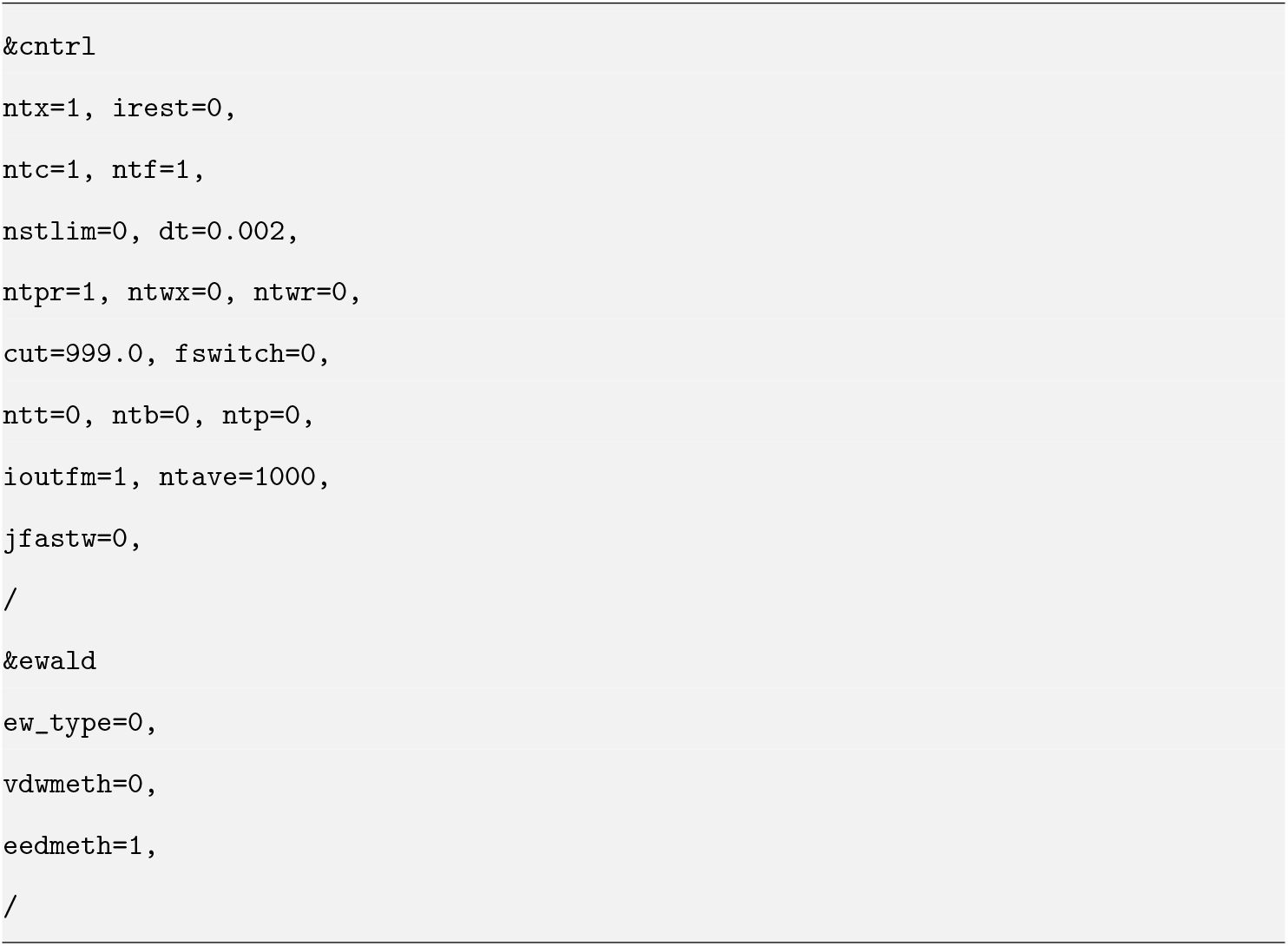

### PSF in NAMD: Single point energy configuration file

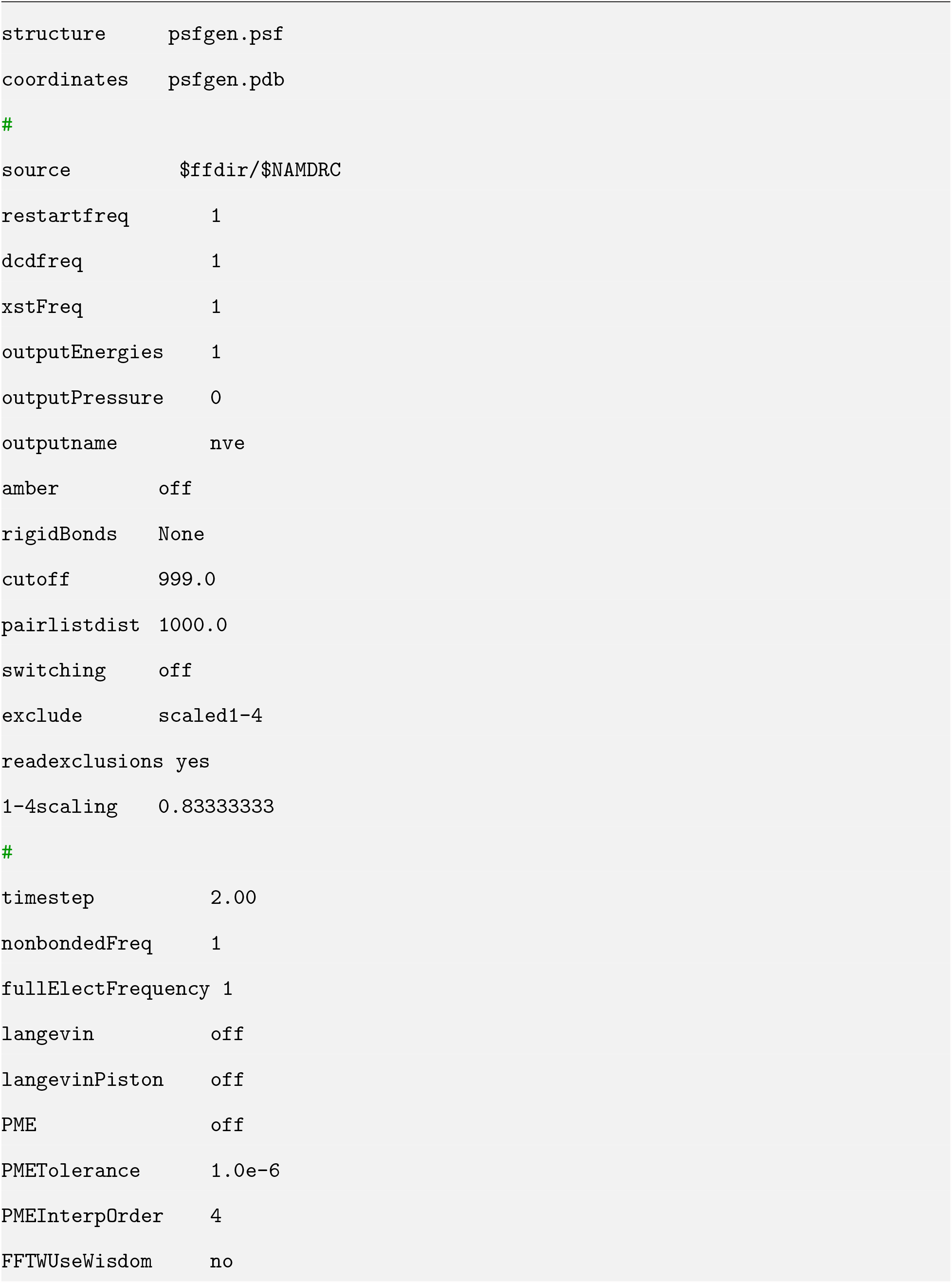
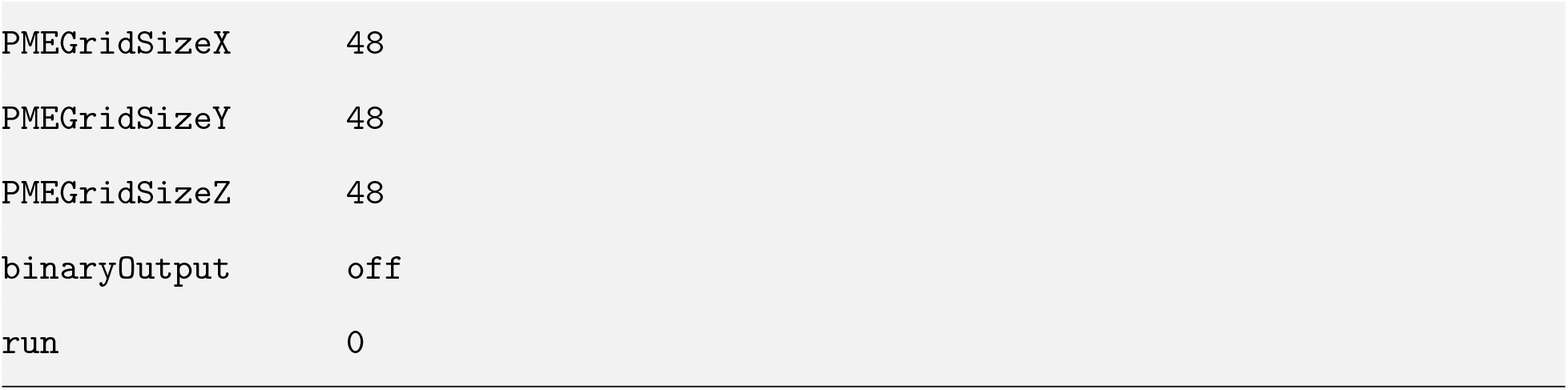

### Protein case study: NAMD configuration file

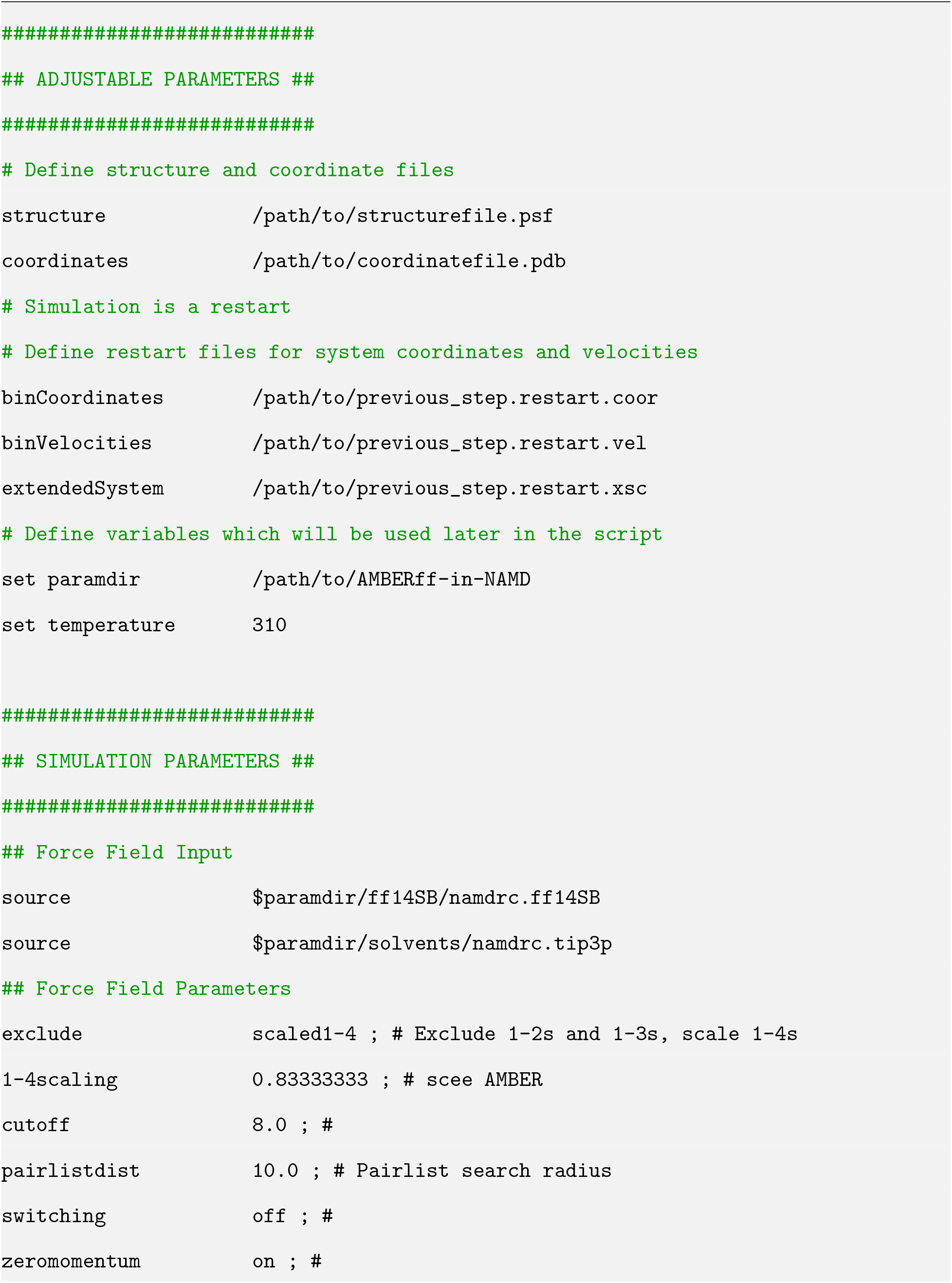
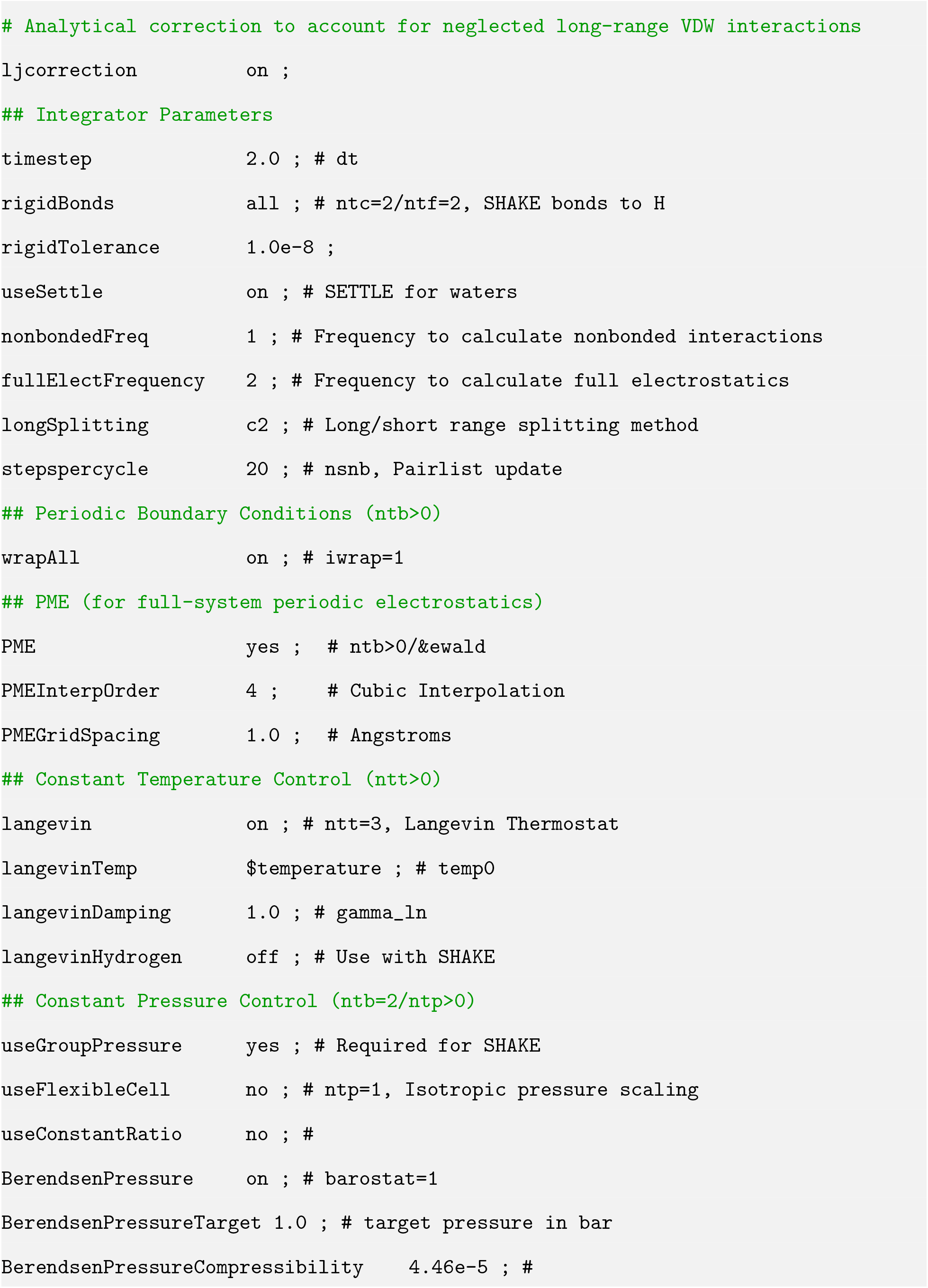
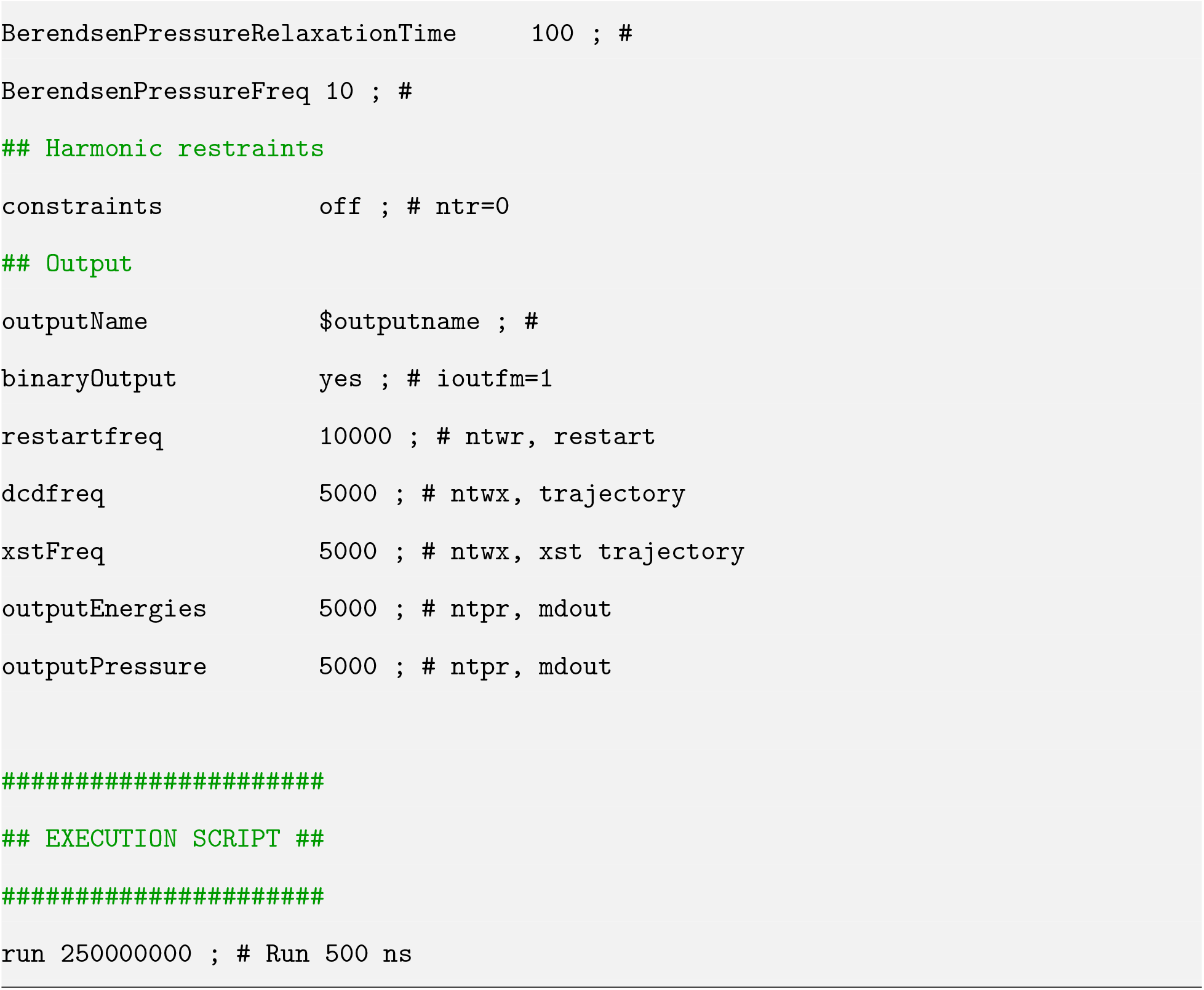

### Lipid case study: NAMD configuration file

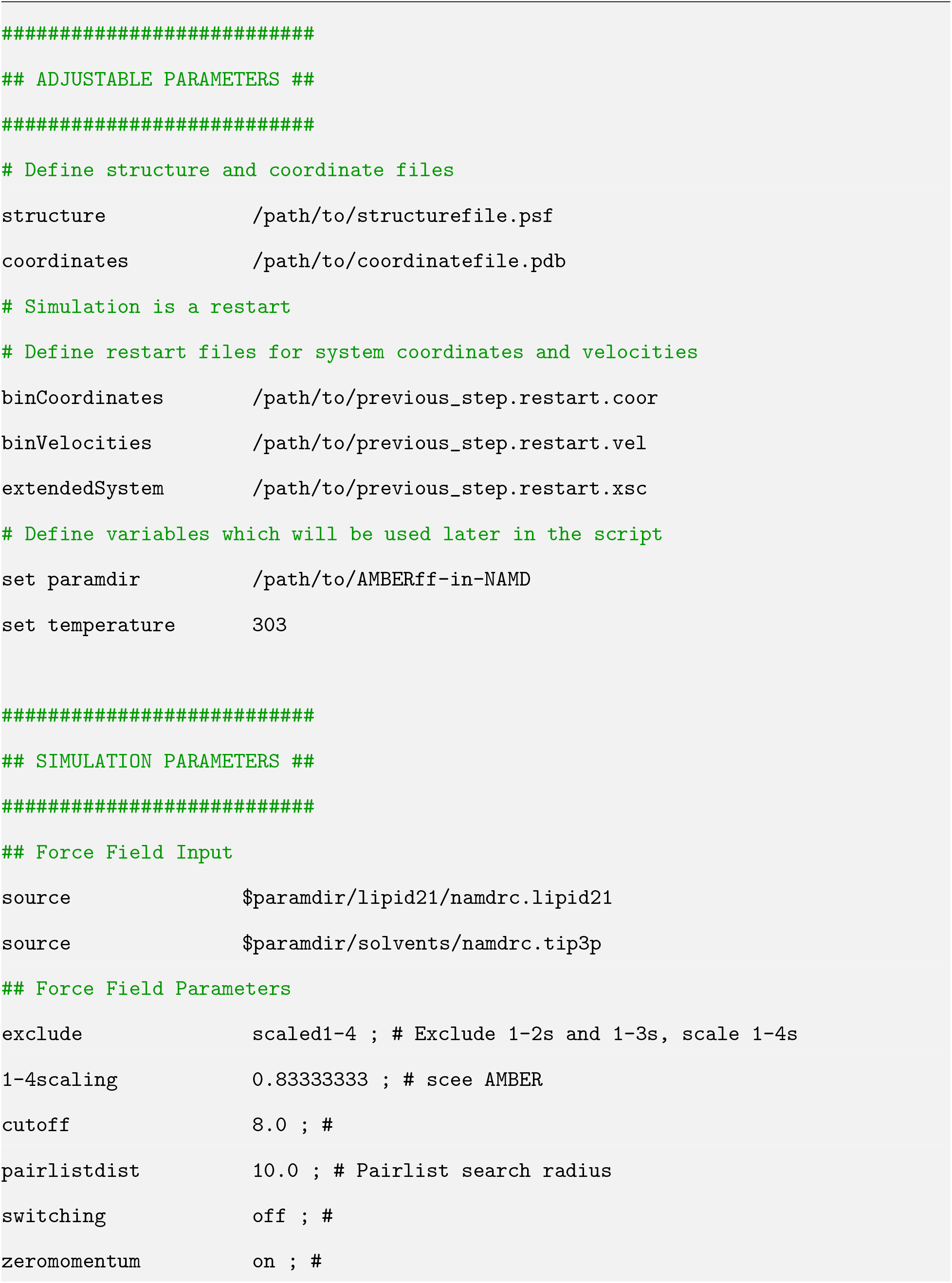
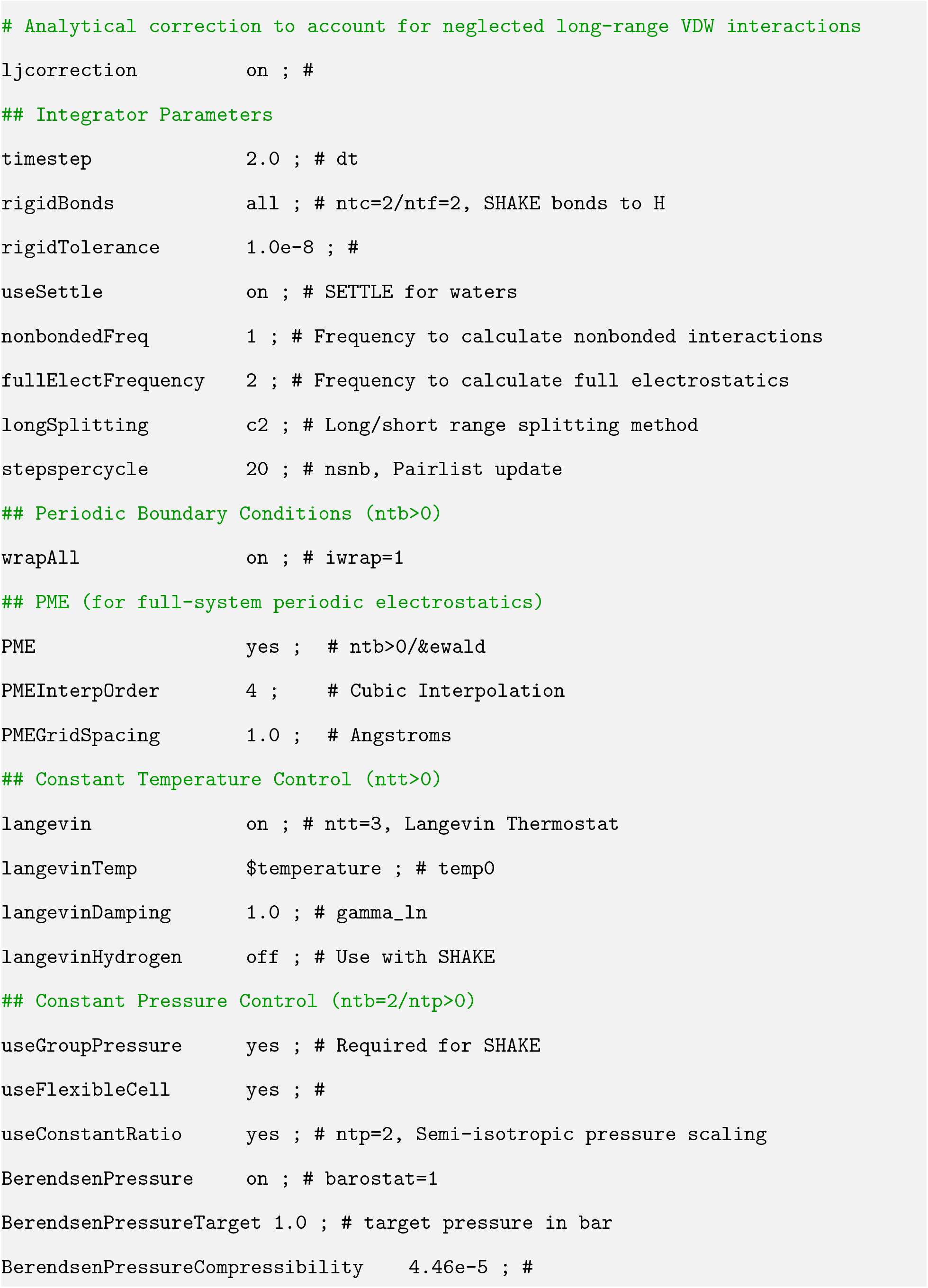
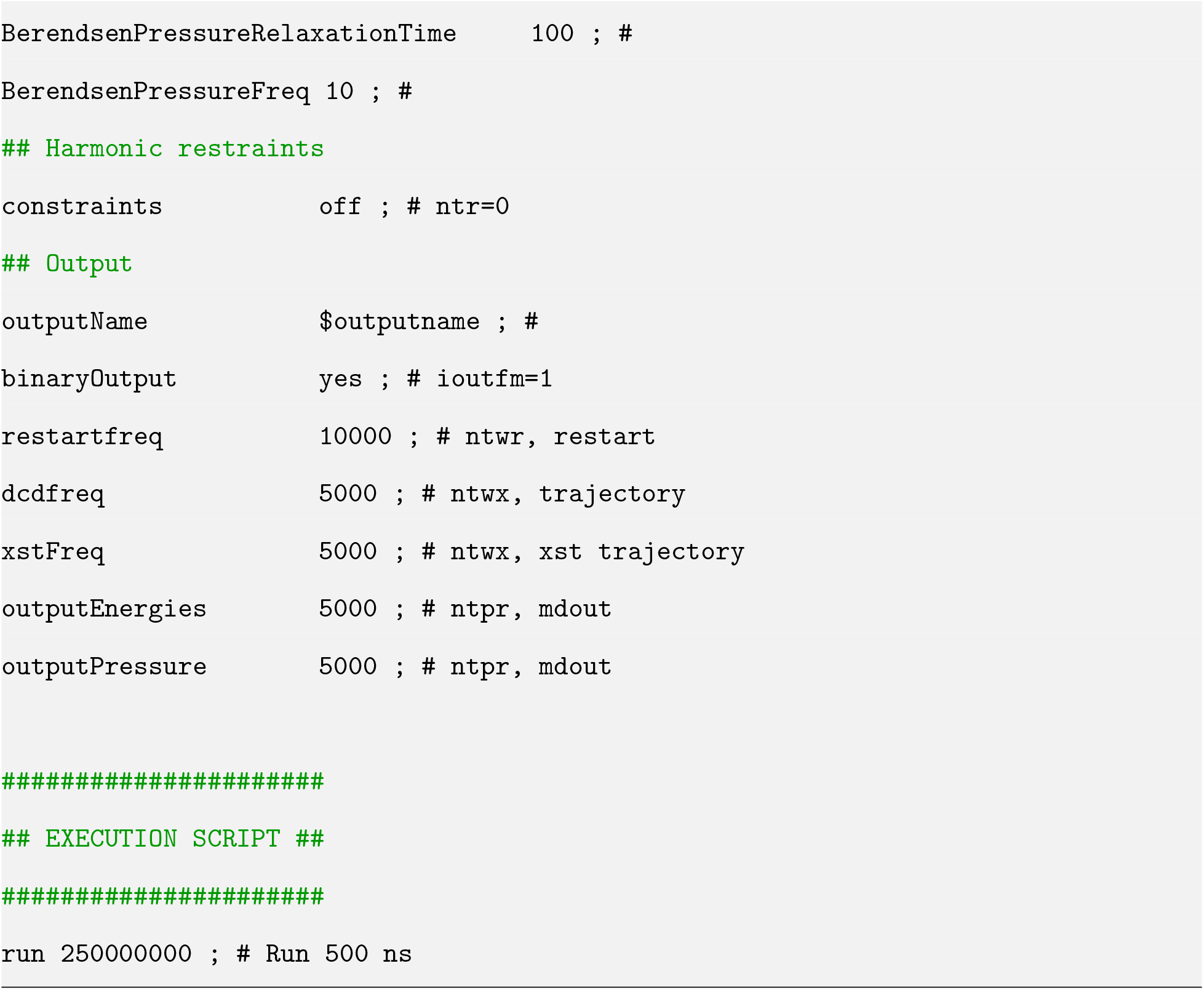

### Nucleic acid case study: NAMD configuration file

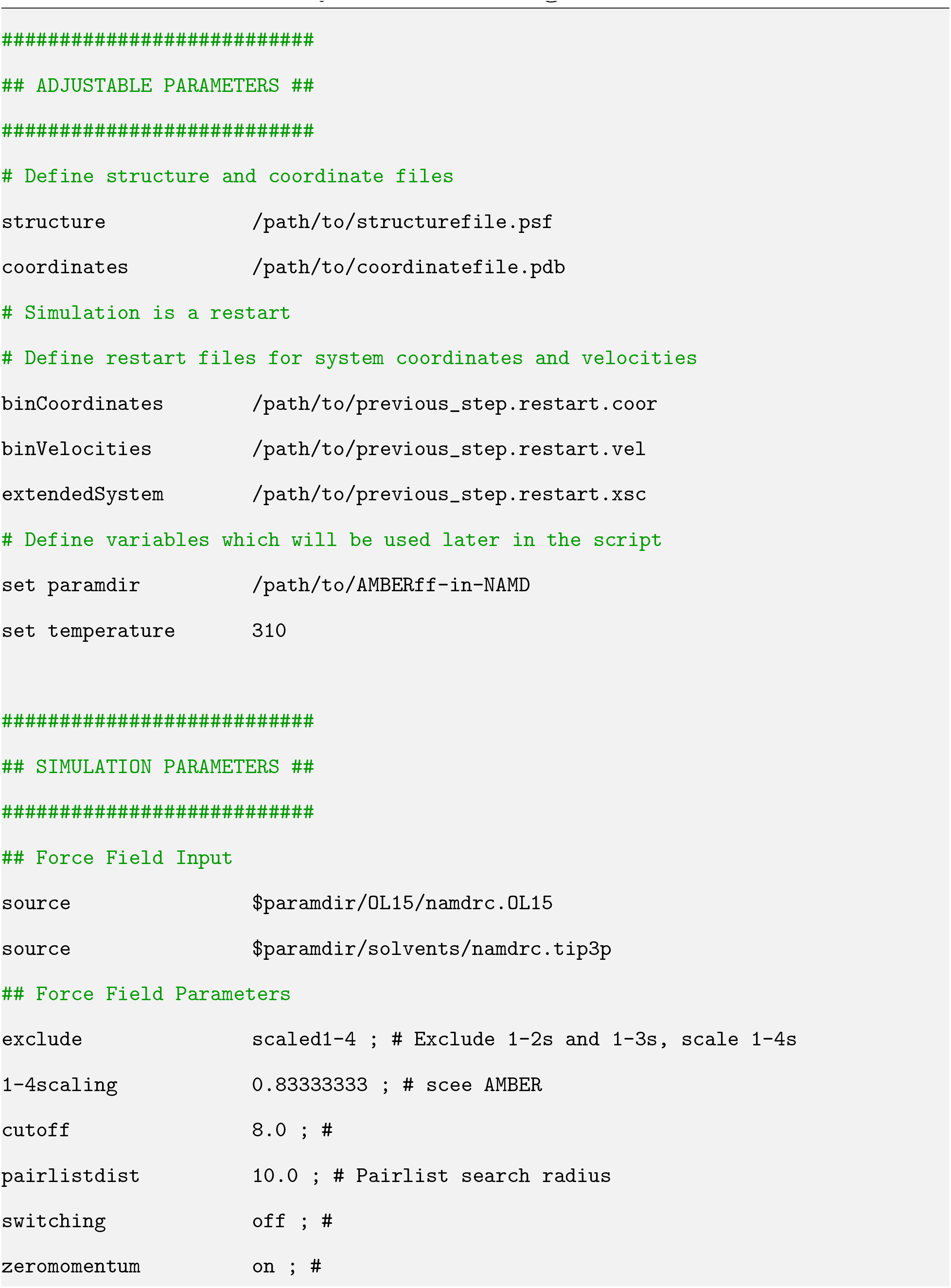
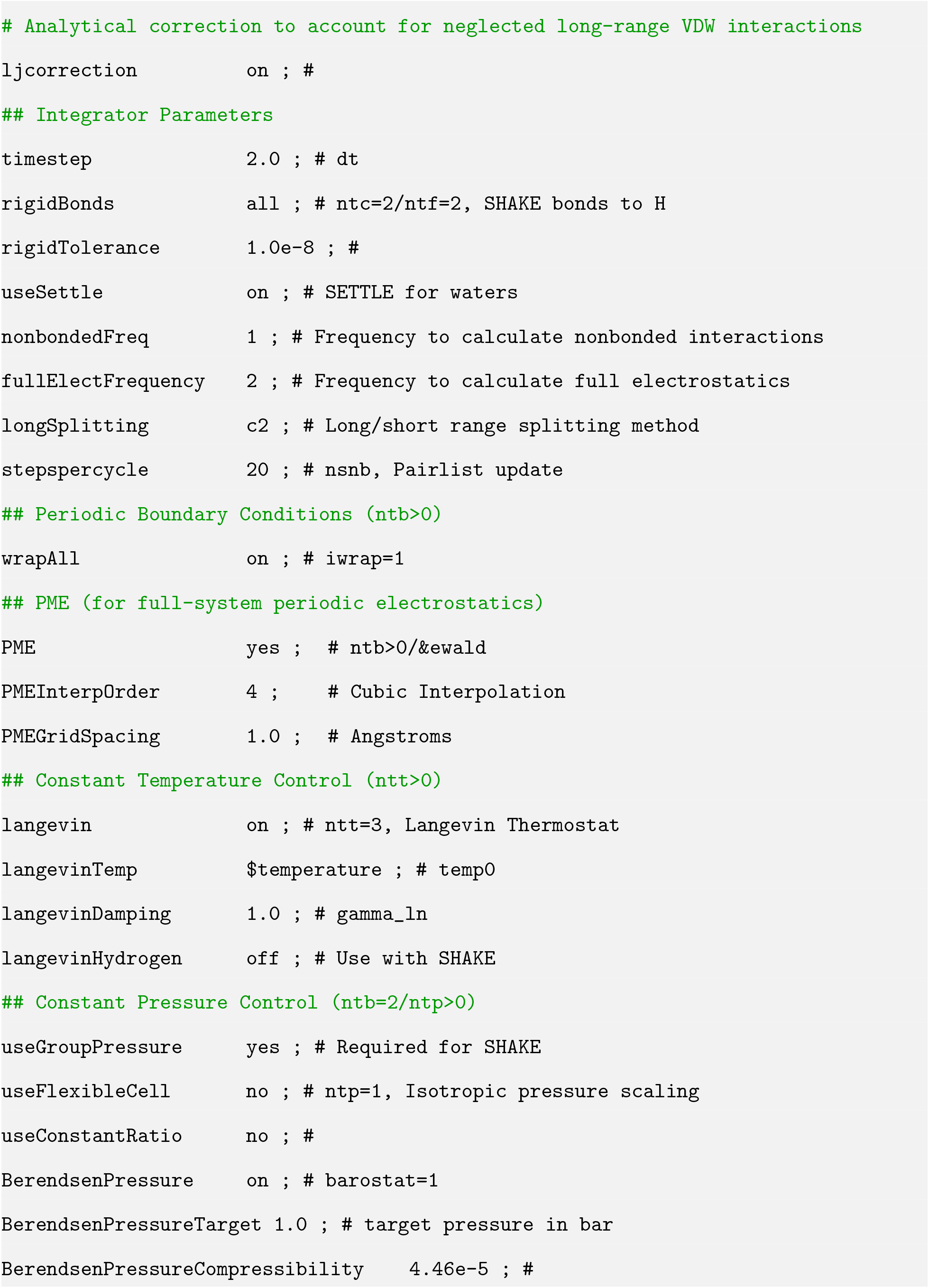
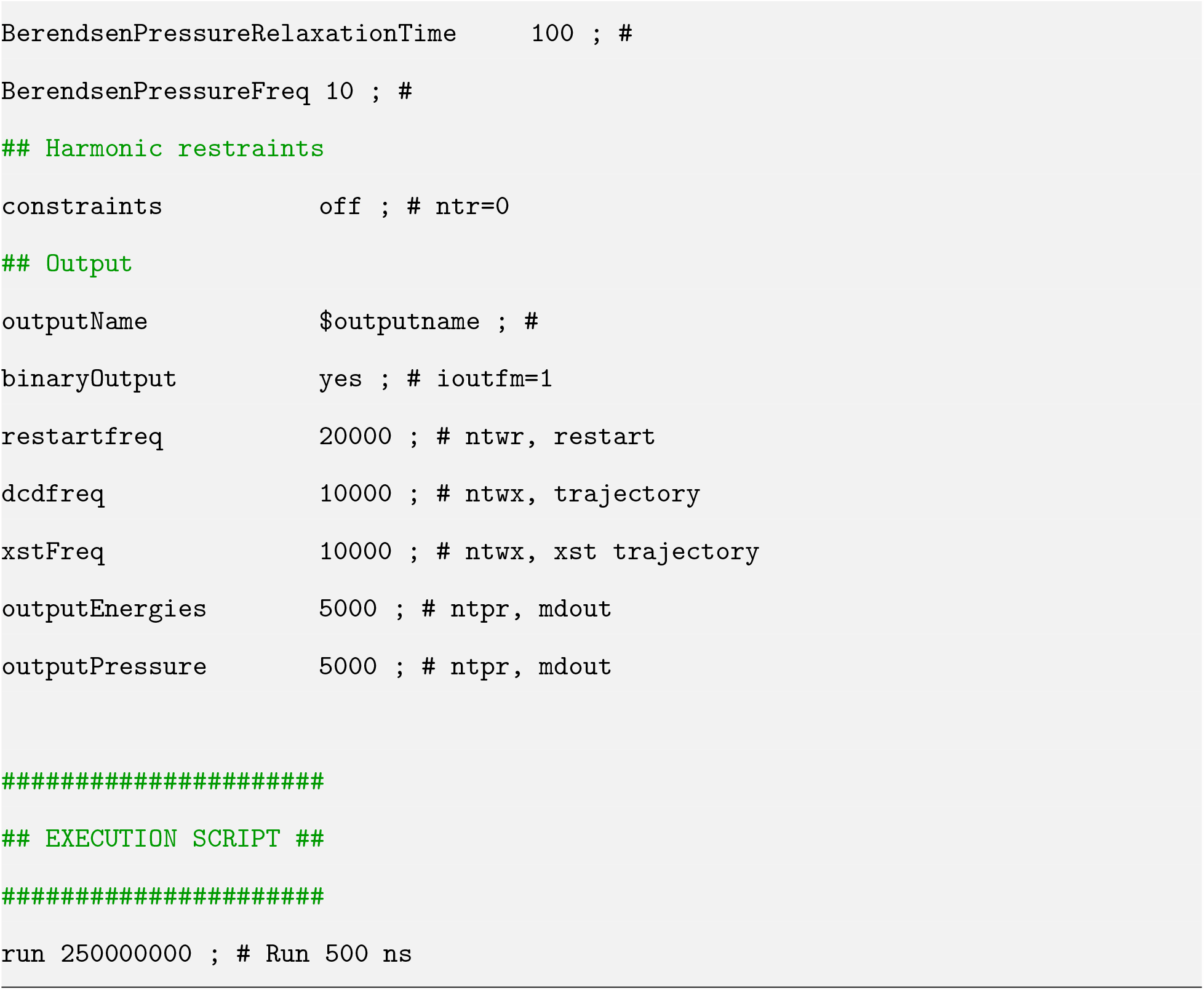

